# A new High-Throughput-Screening-assay for Photoantimicrobials Based on EUCAST Revealed Photoantimicrobials in Cortinariaceae

**DOI:** 10.1101/2021.04.02.438202

**Authors:** Johannes Fiala, Harald Schöbel, Pamela Vrabl, Dorothea Dietrich, Fabian Hammerle, Desirée Josefine Altmann, Ronald Stärz, Ursula Peintner, Bianka Siewert

## Abstract

A spark of light might unravel antimicrobial activity of colored compounds, which otherwise would have been classified as inactive. While many mushrooms contain such colorful pigments but lack antimicrobial activity, we wondered if a controlled irradiation is needed to unleash their effect. To explore such photoantimicrobial actions in the Kingdom Fungi, an efficient high-throughput-screening (HTS) assay is needed. Here we report on the establishment of a reliable photoantimicrobial assay based on the EUCAST recommendations, which was validated with known photosensitzers (i.e., curcumin, phenalenone, rose bengal, an hypericum extract, and methylene blue). Furthermore, an improved LED-irradiation setup enabling with only 24 LEDs a homogenous irradiation of a 96-well plate is presented. The established HTS-assay was utilized to screen six colorful *Cortinarius* extracts unrevealing *C. xanthophyllus* and *C. rufo-olivaceus* as promising sources for new photoantimicrobials.

## 1 Introduction

Whenever microorganisms share the same ecological niche – as for example soil fungi and soil bacteria – an orchestra of chemical compounds evolves reaching from mediators of stimulative symbiosis to detrimental antibiosis (Frey-Klett et al., 2011; Deveau et al., 2018). Plenty of such natural products have commercial values, especially as pharmaceuticals (Hyde et al., 2019). For example, most antibiotics approved by the Food and Drug Administration (FDA) are natural products (Lewis, 2020) and belong to antibiosis, which is described as chemical warfare.

According to Künzler (2018), fungal cells usually defend themselves by rather secreting chemical effectors against microbial competitors, than by storing them intracellular (Künzler, 2018). Nevertheless, fruiting bodies – or more precisely the hyphae differentiating into fruit-body tissues – often contain promising antibiotics. For example, various antimicrobial triterpenoids were isolated from the fruiting bodies of polypores, especially of *Ganoderma spp*. (Dresch et al., 2015; Basnet et al., 2017). These observations are rather the rule than the exception, because for most basidiomycete genera an antimicrobial activity was found in extracts from the fruiting bodies. Just for a few genera, for example *Cortinarius*, antimicrobial activities were infrequently described. This, however, contrasts with the observation that fruiting bodies of this genus are rarely infested by other microorganisms (Moser, 1972). Thus, we were wondering whether an important co-factor was missing in the common screening attempts.

Co-factors, which can influence the antimicrobial activity of a secondary metabolite, might be metals (Lachowicz et al., 2020), pH-conditions (Lee et al., 1997; Wiegand et al., 2015), or just a spark of light (Wozniak and Grinholc, 2018; Dos Santos et al., 2019). Such light-activated defense strategies (Downum, 1992; Flors and Nonell, 2006) are well-known for members of the kingdom Plantae and were recently suggested to be also present in fungi (Siewert and Stuppner, 2019; Siewert et al., 2019; Siewert, 2021). Furthermore, light-activated natural compounds are promising pharmaceuticals (Hudson and Towers, 1991; Berenbaum, 1995; Siewert and Stuppner, 2019).

As part of a putative light-activated defense system, the first photosensitizers, i.e. light-activated chemical compounds, were recently activity-guided discovered in fruiting bodies of macromycota (Siewert et al., 2019; Hammerle et al., 2020). Light-activated antimicrobial effects of basidiomycetes are, however, not described yet, despite promising hints (Siewert, 2021). The lack of described photoantimicrobials might be the consequence of a non-existing photo-antimicrobial high-throughput screening (HTS) assay.

In general, plenty of different antimicrobial susceptibility tests are available determining the minimal inhibitory activity (MIC) of a substance. The utilized techniques reach from diffusion over thin-layer chromatography to dilution methods (Balouiri et al., 2016). In recent years, two standard protocols – one published by the Clinical and Laboratory Standards Institute (Weinstein and Lewis, 2020) and the other by the European Committee on Antimicrobial Susceptibility Testing (EUCAST (Microbiology and Diseases, 2003)) – were established. Most promising for a HTS assays are such microbroth-dilution assays, which are based on visual (CLSI) or spectrophotometric (EUCAST) turbidity measurements (Wiegand et al., 2008). Microbroth-dilution assays can be conducted in 96 well-plates and thus allow a high throughput: Eight antibiotics can be tested in ten different concentrations on one plate in the dark, including the sterility and growth controls (Wiegand et al., 2008).

The crucial part of every PhotoMIC assay is the irradiation. Nowadays, dental curing lights (Nielsen et al., 2015) or handmade LED setups (Morici et al., 2020) replaced previously used light bulbs and lasers (Calin and Parasca, 2009). Dental lights – originally designed to polymerize composite fillings – allow only single irradiation, and therefore limit the throughput. Described LED-setups (not limited to microbials) vary from a single-emitter LED (Ogonowska et al., 2019) over 24 (Quintanar et al., 2016) and 96 LEDs (Butler et al., 2010; Chen et al., 2012; Hopkins et al., 2016; Katz et al., 2018) to 195 (Bajgar et al., 2020) or even 432 diodes (Pieslinger et al., 2006). A drawback of all settings with less than 100 diodes, is the missing homogenous light-distribution throughout a 96-well plate (Chen et al., 2012; Hopkins et al., 2016; Quintanar et al., 2016; Ogonowska et al., 2019). Consequently, only parts of a 96-well plate can be used. Common to all multi-diodes settings is the equidistant arrangement of the diodes along the printed circuit board. Taking the nature of light into account, however, we wondered whether an asymmetric positioning of the diodes might improve the all-over distribution of light. Having extrapolated simulations for single LEDs in mind, we hypothesized that a homogenous illumination with only 24 diodes is possible.

Here we will report on (1) the design of a modular, 24 LEDs based irradiation setup for 96-well plates, (2) the establishment of a HTS-PhotoMic assay which was validated with five standard photosensitizers (PS, curcumin, phenalenone, rose bengal, and hypericin) and five irradiation wavelengths (λ = 428, 478, 523, 598, and 640 nm); And, (3) the results of a sample set existing out of six *Cortinarius* extracts and identifying the basidiomycete *Cortinarius xanthophyllus* and *C. rufo-olivaceus* as species containing photoantimicrobial(s) active against *Staphylococcus aureus* and *Candida albicans*.

## 2 Materials and Methods

### 2.1 Optical simulations, irradiation setup, and light measurements

The irradiation system is based on LED-technology. To achieve uniform irradiance along the entire sample, the arrangement of the individual LEDs within the 6 × 4 LED array is crucial. Therefore, the LED positions were optimized and verified with optical simulations. The simulation is based on an optical model for single LEDs (Wood, 1994) and is modified to calculate irradiance distribution in terms of Cartesian coordinates (Moreno et al., 2006). To simulate the irradiance *E*(*x,y, z*) at any point of the *x, y*-plane at a working distance *z*, the 6 × 4 LED array is modeled as

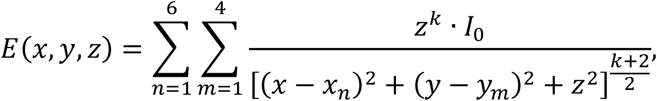

where *x_n_* and *y_m_* are the positions of the individual LEDs in meters and *I*_0_ is the radiant intensity in watt per steradian. The deviation of the manufactured LED from a perfect Lambertian emitter is considered with the correction factor *k*, which depends on the viewing angle *θ*_1/2_

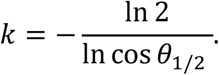

The viewing angle *θ*_1/2_ is the off-axis angle from the LED centerline where the radiant intensity is half of the peak value and is provided by the LED manufacturer. By varying the individual LED positions *x_m_* and *y_m_*, the irradiance distribution in the sample plane can be modified. To achieve a uniform irradiance distribution, the individual LED positions were optimized by a nonlinear least-square curve fitting method with constraints (Betts, 1976; Coleman and Li, 1994; 1996). Optical simulations and optimization were performed using MATLAB R2019b.

All irradiation experiments were carried out with a specially developed irradiation device (SciLED, MCI, Innsbruck) based on LED technology (Figure 1, left). The device consists of an extendable sample holder, where the 96-well plates can be inserted and reproducible positioned in the irradiated area. If the experimental design requires alternative culture plates, e. g. petri dishes, the sample holder can be easily adapted. To ensure a versatile area of application, the device has a modular design. Depending on the demanded irradiation conditions, the LED modular units (Figure 1, insert) can be exchanged. The LED modules were assembled with Luxeon CZ Color Line LEDs (B.V., 2019). Each module consists of 24 LEDs of the same color (nominal peak wavelength). For this work, LEDs of the color violet (λ = 420–430 nm), blue (λ = 465–475 nm), green (λ = 520–540 nm), amber (λ = 585–600 nm), and red (λ = 624–634 nm) were used. The arrangement of the LEDs in the array was optimized to ensure a uniform irradiance. Figure 1 (right) shows the simulation of the irradiance of one LED modular unit. Next to the wavelength, the radiant exposure can be adjusted by a timer and an intensity controller.

**Figure 1:**
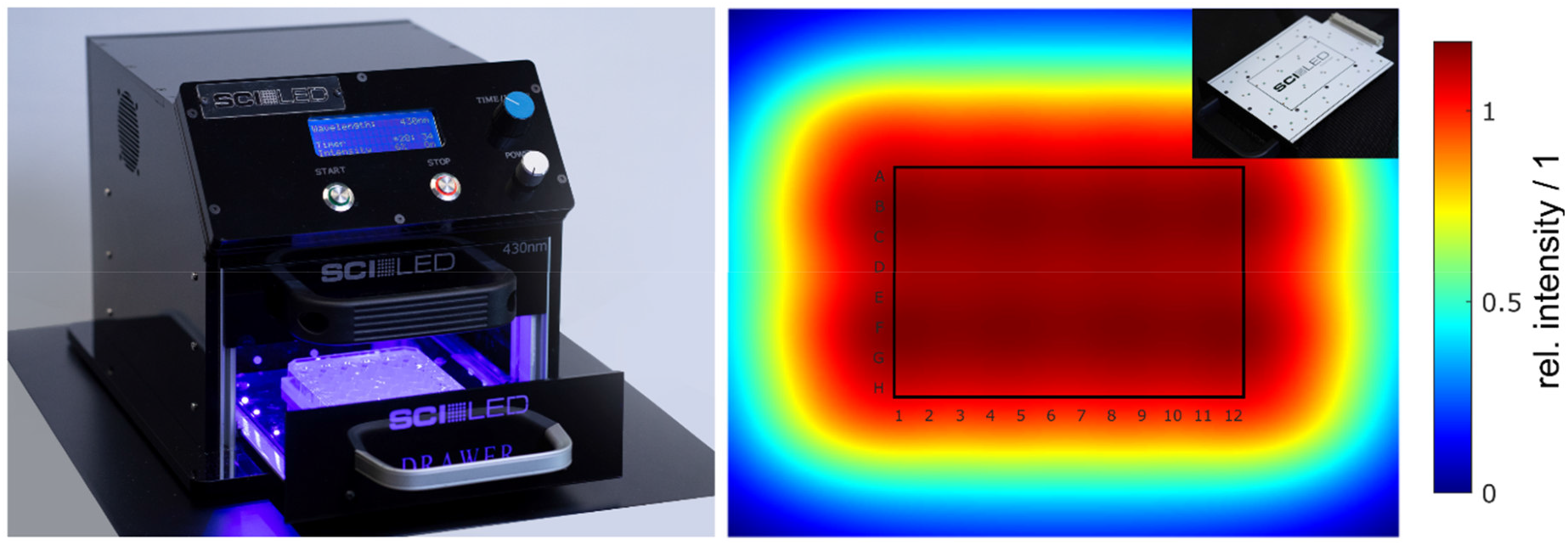
(Color online) Irradiation Setup (SciLED) and simulated irradiance. The irradiation experiments were performed with a LED-based setup (left). Due to its design, the LED modules can be easily exchanged to enable different wavelength settings (insert, top right). With the integrated user interface, the radiant exposure can be set by a timer and an intensity controller. The arrangement of the 6 × 4 LED array on the modules was optimized to achieve a uniform irradiance at the sample plane. Optical simulations of the irradiance show a uniform distribution with a theoretical variation of less than 0.1% over the entire area of a 96-well plates (right).

Light measurements were conducted to characterize the illumination device. To check the uniformity, irradiance was measured using a radiometer and a chemical actinometer (i.e., ferrioxalate). Irradiance measurements were carried out along a 17 mm x 17 mm grid using the radiometer PM100D and the photodiode power sensor S120 VC with a measurement uncertainty of ±3% (λ = 440 – 980 nm) and +5% (λ = 280 – 439 nm) (Thorlabs). The ferrioxalate actinometer (K_3_[Fe(C_2_O_3_)_3_]) and phenanthroline-based developing solutions were made using previously published methods (Hopkins et al., 2016). Spectral measurements were performed using the spectrometer MAYA 2000 Pro equipped with diffraction grating #HC-1 and entrance slit of 5 μm (Ocean Insights), resulting in a spectral resolution of 0.66 nm FWHM. Light was coupled into the spectrometer via an optical fiber with a core diameter of 600 μm (QP600-1-SR-BX, Ocean Insights) and a cosine corrector (CC-3-UV-S, Ocean Insights). The spectrometer was calibrated with a wavelength calibration source (mercury-argon HG-2, Ocean Insights). To characterize the spectral power distribution, the peak shape was modeled with a sum of Gaussian functions (Reifegerste and Lienig, 2008; Supronowicz and Fryc, 2019). By fitting the sum of Gaussian functions to the spectral data, the wavelength where the intensity maximum occurs (peak wavelength in nm), the full width at half of the intensity maximum (FWHM in nm), and the full width at ten percent of the intensity maximum (FW 0.1 · *I_max_* in nm) were calculated.

To evaluate the uniformity, the arithmetic mean irradiance *E_m_*, the standard deviation *SD* and the coefficient of variation *cv* were calculated. As the uniformity was simulated and measured using a radiometer and a chemical actinometer, a comparison with the dimensionless parameter *cv* is convincing. The coefficient of variation is calculated as the ratio of the standard deviation and the arithmetic mean. Ensuring a precise representation of the spectral data by the model with Gaussian functions, the fit was accepted with a coefficient of determination *R*^2^ larger than 0.999.

### 2.2 Mycochemical Part – Reagents, Instruments, and Methods

All solvents for the extraction and isolation processes were purchased from VWR International (Vienna, Austria). Acetone was distilled prior to use. Solvents for HPLC experiments had pro analysis (p.a.) quality and were obtained from Merck (Merck KGaA, Darmstadt, Germany). Ultrapure water was obtained with the Sartorius arium® 611 UV purification system (Sartorius AG, Göttingen, Germany).

Desiccation of the collected fungi was achieved with a dörrex® drying-apparatus from Stöckli (A. & J. Stöckli AG, Switzerland) operated at a temperature of 50 °C. The fungal biomaterial was milled with a Bosch rotating coffee grinder MKM 6003 (Stuttgart, Germany). The samples were weight with scales from KERN ALS 220-4 (KERN & SOHN GmbH, Balingen-Frommern, Germany) and Sartorius Cubis®-series (Sartorius AG, Göttingen, Germany). During the extraction process, the ultrasonic bathes Sonorex RK 106, Sonorex RK 52, and Sonorex TK 52 (BANDELIN electronic GmbH & Co. KG, Berlin, Germany) were utilized. Vortexing was done with a Vortex-Genie 2 mixer (Scientific Industries, Inc., Bohemia, New York). For centrifugation, an Eppendorf 5804R centrifuge with a F-45-30-11 – 30 place fixed angle rotor (Hamburg, Germany) was used.

HPLC measurements were carried out with the modular system Agilent Technologies 1260 Infinity II with a quaternary pump, vial sampler, column thermostat, diode-array detector, and mass spectrometer. Moreover, the HPLC-system Agilent Technologies 1200 Series with a binary pump, autosampler, column thermostat, and diode-array detector was used. HPLC-systems were purchased from Agilent Technologies, Inc. (Santa Clara, USA). For all HPLC measurements, a Synergi 4u MAX-RP 80A 150 x 4,60mm column was used. HPLC-DAD-ESI-MS analysis was carried out with the modular system Agilent Technologies 1260 Infinity II equipped with a quaternary pump, vial sampler, column thermostat, diode-array detector, and an ion trap mass spectrometer (amaZon, Bruker, Bremen, Germany).

Pipetting was done with pipettes and tips from Eppendorf AG (Hamburg, Germany) and STARLAB International GmbH (Hamburg, Germany). Reagent reservoirs were obtained from Thermo Fischer Scientific (Waltham, Massachusetts, USA).

### 2.3 Mycochemical Part

#### 2.3.1 Preparation of fungal extracts

The fungal biomaterial was dried on a desiccator (T ~ 50°C) right after collection (see Table S1) and stored at room temperature until further use (T = 23.0°C, humidity = 20 +/− 5%). The biomaterials were milled and sieved utilizing a mesh with the size of 400 μm. The extraction process was performed under light exclusion at room temperature. The powdered materials (m = 2.00 g) were extracted with acidified acetone (V = 20 ml, 0.1 v/v% 2N HCl) in an ultrasonic bath (t = 10 min). After centrifugation (t = 10 min, T = 4°C, F = 20817 g), acetone was decanted and filtered through cotton wool. The fungal material was extracted twice more with acidified acetone (V = 5 ml). After centrifugation, the supernatant was collected, evaporated, and stored in brown glass vials at room temperature (see Table 1 for yields).

**Table 1:** Optical characterization of the irradiation device. The spectral power distributions and the irradiance at the sample plane were measured for all LED module colors. From the spectral data, the actual peak wavelength, the full width at half of the intensity maximum (FWHM), and the full width at ten percent of the intensity maximum (FW 0.1 · *I_max_*) were obtained by fitting a sum of Gaussian functions to the data. From the irradiance measurements, the arithmetic mean (*E_m_*), the standard deviation (*SD*) and the coefficient of variation (*cv*) were calculated.

| LED module<br>color | spectral information |  |  |  | irradiance |  |  |
| --- | --- | --- | --- | --- | --- | --- | --- |
| | wavelength | FWHM | FW 0,1 · $I_{max}$ | $R^2$ | $E_m$ | $SD$ | $cv$ |
|  | [nm] | [nm] | [nm] | [1] | [mW/cm<sup>2</sup>] | [mW/cm<sup>2</sup>] | [1] |
| violet | 428 | 15 | 36 | 0.9994 | 13 | 1.0 | 0.081 |
| blue | 478 | 27 | 63 | 0.9991 | 8.7 | 0.70 | 0.076 |
| green | 523 | 33 | 78 | 0.9998 | 6.0 | 0.44 | 0.073 |
| amber | 598 | 16 | 38 | 0.9994 | 1.1 | 0.084 | 0.078 |
| red | 640 | 18 | 45 | 0.9999 | 6.4 | 0.47 | 0.074 |

#### 2.3.2 Reagents, Instruments, and Methods

Curcumin, dimethylsulfoxid (DMSO), lysogeny broth (LB) agar, phenalenone, and RPMI1640 medium were received from Merck KGaA (Darmstadt, Germany). Potato dextrose agar (PDA) and Mueller Hinton Broth (MHB) were purchased from VWR International (Vienna, Austria). Rose Bengal (RB) was received from TCI Europe (Zwijndrecht, Belgium). *Hypericum perforatum* extract (ethanol) was prepared from the pharmaceutical drug “Johanniskraut 600 mg forte” by Apomedica (Graz, Austria). The 96-well plates (flat bottom) were bought from Sarstedt (Nümbrecht, Germany).

The U-2001 spectrophotometer for adjusting the McFarland standard was from Hitachi (Chiyoda, Japan). For measurement of the 96-well plates, a Tecan Sunrise Remote Plate Reader was used (Männedorf, Switzerland). The adjustment of pH-values was carried out with the pH-meter Mettler Toledo SevenMulti (Mettler-Toledo GmbH, Vienna, Austria).

Pipetting was done with pipettes and tips from Eppendorf AG (Hamburg, Germany) and STARLAB International GmbH (Hamburg, Germany). Reagent reservoirs were obtained from Thermo Fischer Scientific (Waltham, Massachusetts, USA).

#### 2.3.3 Strains and Cultivation

All experiments on photodynamic inhibition (PDI) of growth of microorganisms (MOs) and the preparations were carried out under aseptic conditions in a laminar airflow cabinet at room temperature. The test strains used in this study were *Candida albicans* (501670), *Escherichia coli* (DSM1103), and *Staphylococcus aureus* (DSM1104). The strains were reactivated from frozen state and prepared according to manufacturer’s recommendations (https://www.dsmz.de/). Until further use, bacterial cultures were stored in darkness at 4°C on lysogeny broth agar. *C. albicans* was cultivated on potato dextrose agar under the same conditions.

For the PDI experiments, the stored cultures were reactivated, and an overnight culture was incubated (T = 37°C, t = 24 h, dark conditions). The bacterial culture inoculum was prepared using a spectrophotometer measurement at λ = 600 nm. Turbidity was adjusted to a McFarland standard of 0.5 to prepare a standard suspension of 1.5×10^8^ colony forming units (CFU)/ml. For yeast suspensions, turbidity was measured at λ = 530 nm. Liquid media used for PDI experiments were MHB for bacteria and RPMI-1640 (double strength) for yeast.

#### 2.3.4 PhotoMIC Assay

For the PDI experiments, flat-bottom 96-well plates were used. On each plate, an extract test section, growth control, fraction-blank, medium-blank, and sterility controls were set up (Figure 2). In the test section three concentrations of fungal extracts (i.e., c = 25 μg/mL, 50 μg/mL, and 75 μg/mL), were tested. If needed, smaller or larger concentrations were tested as well. Further, positive controls (dark condition) were established for each experiment: Curcumin (c = 30 μg/mL, 81.5 μM) for *C. albicans*, phenalenone (c = 75 μg/mL, 416.2 μM) for *E. coli*, and phenalenone (c = 25 μg/mL, 138.7 μM) for *S. aureus*.

**Figure 2.**
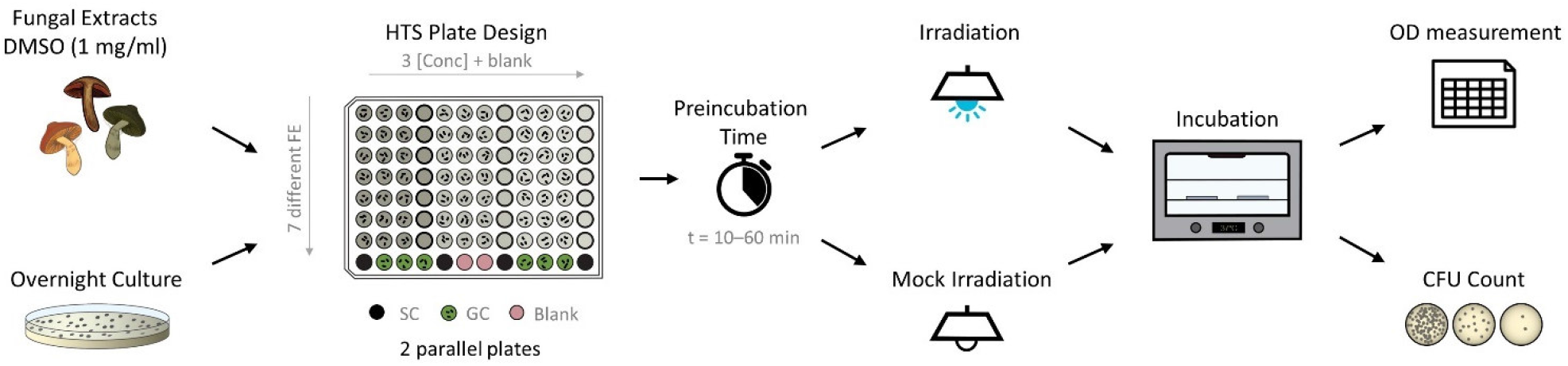
Flow chart of photo-antimicrobial HTS based on the microdilution method. The pipetting scheme represents the fast screening approach of extracts. Up to seven different fungal extracts with three concentrations each are tested on one plate. SC… sterility control, GC… growth control

Positive controls, fungal extracts (FE), and growth control were inoculated with an inoculum (V = 50 μl) of the test strains within t = 30 min after the turbidity adjustment. Two identical 96-well plates were prepared for both dark and light treatment. After ten or sixty minutes of preincubation time in darkness, one plate was irradiated with the SciLED panel at λ = 470 nm for t = 19 min 8 sec, corresponding to a light dose of *H* = 9.3 J/cm^2^. An alternative irradiation setup was t = 61 min 44 sec, corresponding to a light dose of *H* = 30 J/cm^2^. The other plate was kept in the darkness at room temperature.

After (mock)irradiation, the plates were submitted to turbidity measurements. Here, a plate reader was used and before measuring the optical density (bacteria: λ = 600 nm, fungi: λ = 530 nm), the plates were shaken for five seconds. Viability controls were drawn from the control vials and plated on LB/PDA agar. Afterwards, the 96-well plates and LB/PDA agar plates were incubated at T = 37°C in the dark for 24 hours. A second measurement of turbidity was done, followed by taking samples of wells that showed inhibition (>20%) of population growth control.

Assessment of the PDI experiment was done by correlating the treated well to the uninhibited growth control. Turbidity of fraction-blank and medium-blank was subtracted from corresponding wells to eliminate deviation caused by darkening or bleaching of media and extracts. Each concentration of fungal extracts, the positive control, and the growth control were measured at least in triplicates.

### 2.4 Singlet-Oxygen Detection via the DMA-Assay

To analyze the ability of the six fungal extracts (FE) to generate singlet oxygen after irradiation, the previously described dimethyl anthracene (DMA) assay and a previously characterized irradiation setup were employed (Siewert et al., 2019). As a first step, a DMA solution in ethanol (c = 1.4 mM) (L1) and a L-ascorbic acid-solution in ultrapure water (c = 100 mM, pH = 7.0-7.4) (L2) were prepared. The fungal extracts were dissolved in DMSO (c = 1 mg/mL, FE) and subsequently mixed with the stock solutions (L1 and L2) as well as pure ethanol (L3) to obtain four test-solutions (V = 10 μL FE + 190 μL test-solution): (1) a pure ethanolic solution of the FE to observe photochemical changes of the extract due to the irradiation, (2) a mix with DMA to detect singlet oxygen, (3) a mix with DMA and the antioxidant L-ascorbic acid to prove that singlet oxygen caused the oxidation of DMA, and (4) a control consisting of an ethanolic solution of the extract and L-ascorbic acid to control, that no undesired reaction occurs. DMSO (V = 10 μL) was used as negative control, berberine (c = 1 mg/mL, 2.97 mM, DMSO, V = 10 μL) and RB (c = 0.1 mg/mL, 0.10 mM, DMSO, V = 10 μL) were used as positive controls. Thereafter, optical densities at the wavelengths λ = 377 nm, 468 nm, and 519 nm were measured with a plate reader (t = 0 min), followed by four cycles of blue light (λ = 468 nm, 1.24 J cm^−2^ min^−1^, berberine = positive control) or of green light irradiation (λ = 519 nm, 0.92 J cm^−2^ min^−1^, rose bengal = positive control). All measurements were done as technical duplicates. The results of the DMA-assay were presented as the mean ± standard error. Differences between the relative singlet oxygen formation values were statistically evaluated using one-way ANOVA followed by the Bonferroni post-test, and p < 0.05 was considered to the significant.

## 3 Results

### 3.1 Uniform irradiance and irradiation conditions

The nonlinear optimization of the individual LED positions in the array resulted in a symmetric but not equidistant arrangement. The objective was to achieve a homogenous irradiation distribution with theoretical variations below five percent in an irradiated area of 120 mm × 90 mm (*x* × *y*), which approximately corresponds to the size of a 96-well plate. After several optimization steps, a calculated coefficient of variation *cv* = 0.08% was achieved in the optical simulations. Such uniformity was obtained by decreasing the relative spacing between the outside LEDs and positioning them beyond the area of the irradiated sample. The individual positions of the 24 LEDs are shown in Figure 3. Experimental evaluation of the uniformity resulted in an actual variation between *cv* = 7% and *cv* = 8% for the irradiance measurements and a variation of *cv* = 9% for the chemical actinometer measurements. Over the entire area of a 96-well plate, the resulting irradiance distribution is homogeneous, and from the uniformity standpoint, all 96 wells can be used for irradiation tests. Results of the optimization and irradiance distribution within the 96-well plates are shown in Figure 3.

**Figure 3:**
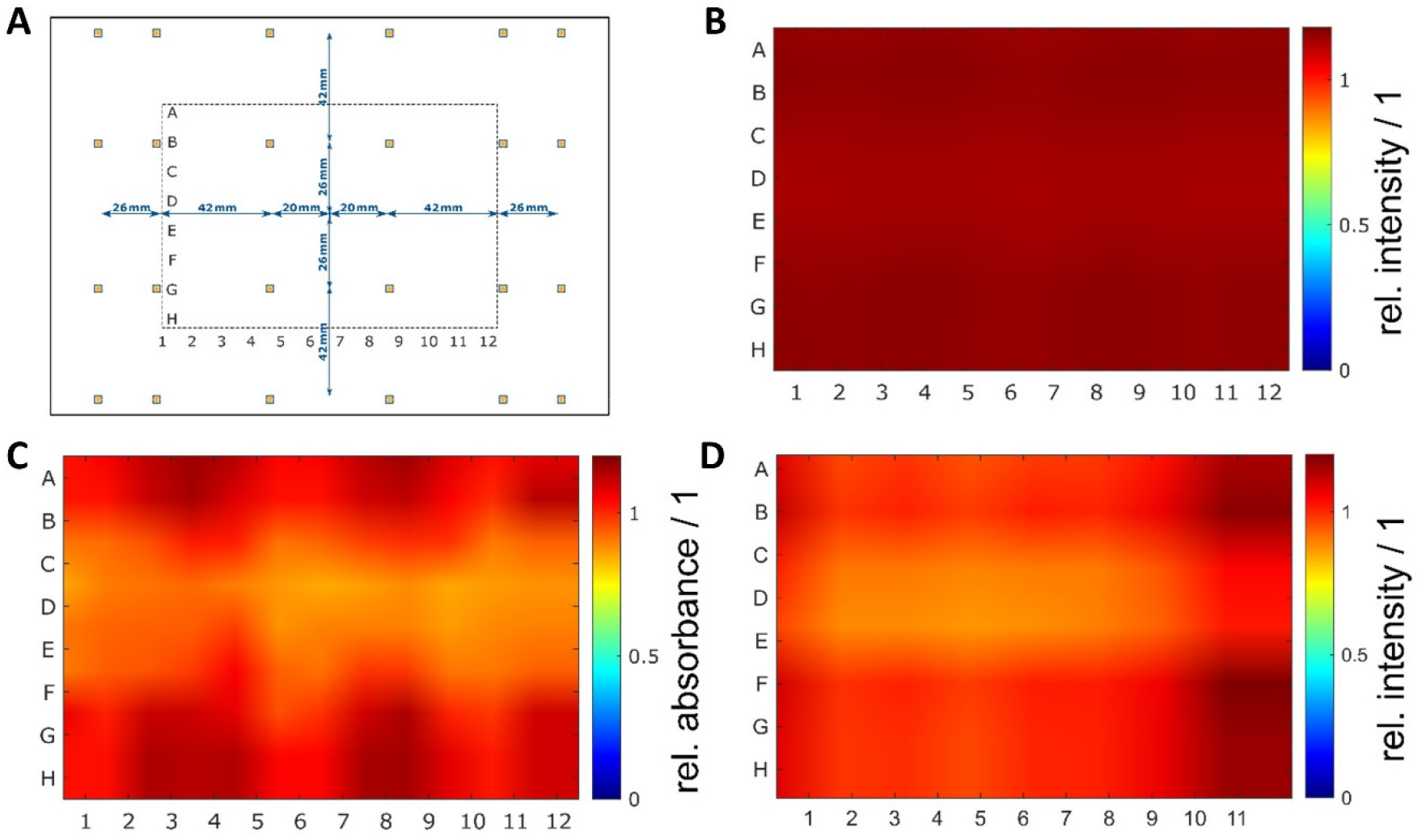
(Color online) Non-equidistant LED arrangement and results of homogeneity measurements. A nonlinear optimization of the individual LED positions in the ***xy***-plane of the printed circuit board resulted in in a symmetric but not equidistant arrangement with decreasing distances on the outside. For a better understanding of the LED arrangement, the position of the irradiated plate is illustrated as well (Figure 2a). The irradiance distribution at the sample plane was determined by optical simulations (Figure 2b) and measured with a chemical actinometer (Figure 2c) and a radiometer (Figure 2d). In the optical simulations a very high uniformity with a theoretical coefficient of variation less than 0.1% (*cv_sim_* = 0.08%) were calculated. Experimental evaluation of the uniformity resulted in variations of less than 10% (chemical actinometer *cv_act_* = 9% and radiometer *cv_rad_* = 8%).

To fully characterize the irradiation device, the spectral power distribution and irradiance were measured for every LED module (violet, blue, green, amber, and red). From these measurements, several spectral parameters, including the actual peak wavelength and the full width at half maximum, were calculated and the average irradiance was determined. Spectral power distributions varied between 15 nm FWHM for violet LEDs and 33 nm FWHM for green LEDs. Average irradiance was the highest for the violet LED module with *E_m_* = 13 + 1.0 mW/cm^2^ and the lowest for the amber LED module with *E_m_* = 1.1 + 0.08 mWW. All results on the irradiation conditions are reported in Table 1.

### 3.2 Establishment of a HTS-Protocol

A high-throughput assay was developed based on the gold-standard microdilution method (Benkova et al.; Wiegand et al., 2008). Like the classic method, the HTS started with an overnight culture of the selected test organisms (*E. coli, S. aureus*, and *C. albicans*) and, separately, with the test compounds or extracts of interest (Figure 2). In the next step, a stock solution of the test extracts or compounds was generated in DMSO and successively diluted in media. MHB was used for the bacteria, while the yeast was cultured in RPMI, double strength. In Figure 2, a flow chart is displayed, including the pipetting scheme for the testing extracts. In Figure S1, the respective flow chart with a pipetting plan for pure compounds is shown. In contrast to the classic microdilution assay, a blank of each tested compound was needed to avoid false-negative effects in the final OD reading which determines the MIC. The next step was a preincubation step, followed by an irradiation step with the chosen wavelength and light doses. A dark control was conducted in parallel to examine the effect of light. After the (mock)-light treatment step, the plates were incubated for t = 24h. Finally, an OD measurement was performed to quantify the MIC, and -if needed- the treated dilutions were submitted to a CFU count to determine the MBC.

### 3.3 Establishment of the Photoantimicrobial Assay and its Validation with known PSs

In the first step, the light tolerance of the test organisms (i.e., *E. coli, S. aureus*, and *C. albicans*) was examined. To achieve this, the microorganisms were irradiated utilizing the five different LED-modules with light doses up to *H* = 30 J/cm^2^. Under all tested light conditions (Figure 4), the irradiated populations were not affected compared to the non-irradiated control groups. Therefore, all observed effects will be due to a combined effect of the light and the test compound/extract.

**Figure 4:**
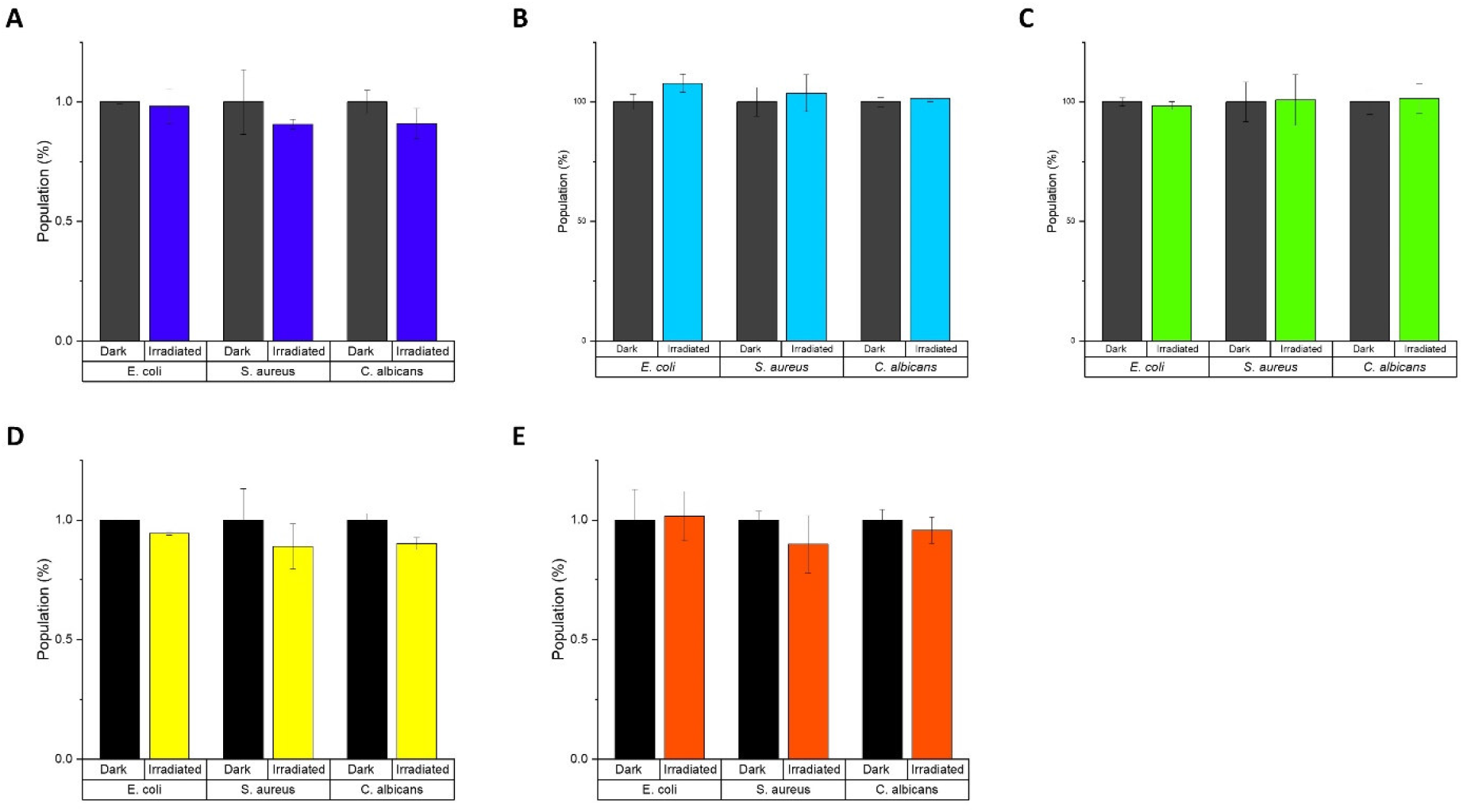
Effect of light irradiation on the growth of *E. coli, S. aureus*, and *C. albicans*. A) λ_irr_ = 428 nm, *H* = 30 J/cm^2^, B) λ_irr_ = 478 nm, *H* = 30 J/cm^2^, C) λ_irr_ = 523 nm, *H* = 30 J/cm^2^, D) λ_irr_ = 598 nm, H= 9.3 J/cm^2^, and E) λ_irr_ = 640 nm, *H* = 30 J/cm^2^.

Next, well-established photosensitizers were selected. In detail, phenalenone, curcumin, rose bengal (RB), methylene blue (MB), and a *Hypericum perforatum* (HP) extract (photoactive ingredient: hypericin) were chosen to validate the irradiation setup. These positive controls (PCs) were characterized by absorption properties complementary to the LED-emission profiles (Figure 5). As depicted, several LED-modules can activate individual PCs, as their absorbance bands fit more than one LED-module. In Table 2, the PhotoMIC values – generated in accordance with the EUCAST guidelines – are given. For each LED-module and tested microorganism, the most active PS is represented in Figure 5, though for several LED-modules a selection of PSs worked. For example, the growth of *S. aureus* was not only impeded with yellow light (λ_irr_ = 523 nm, 30 J/cm^2^) and RB (c = 6 μg/mL, Table 2), but also with yellow light (λ_irr_ = 523 nm, 30 J/cm^2^) and HP (c = 150 μg/mL). The MIC using RB (c = 6 μg/mL), however, was more promising and is thus displayed in Table 2. The doses-response curves are depicted in Figure S2-S6.

**Figure 5.**
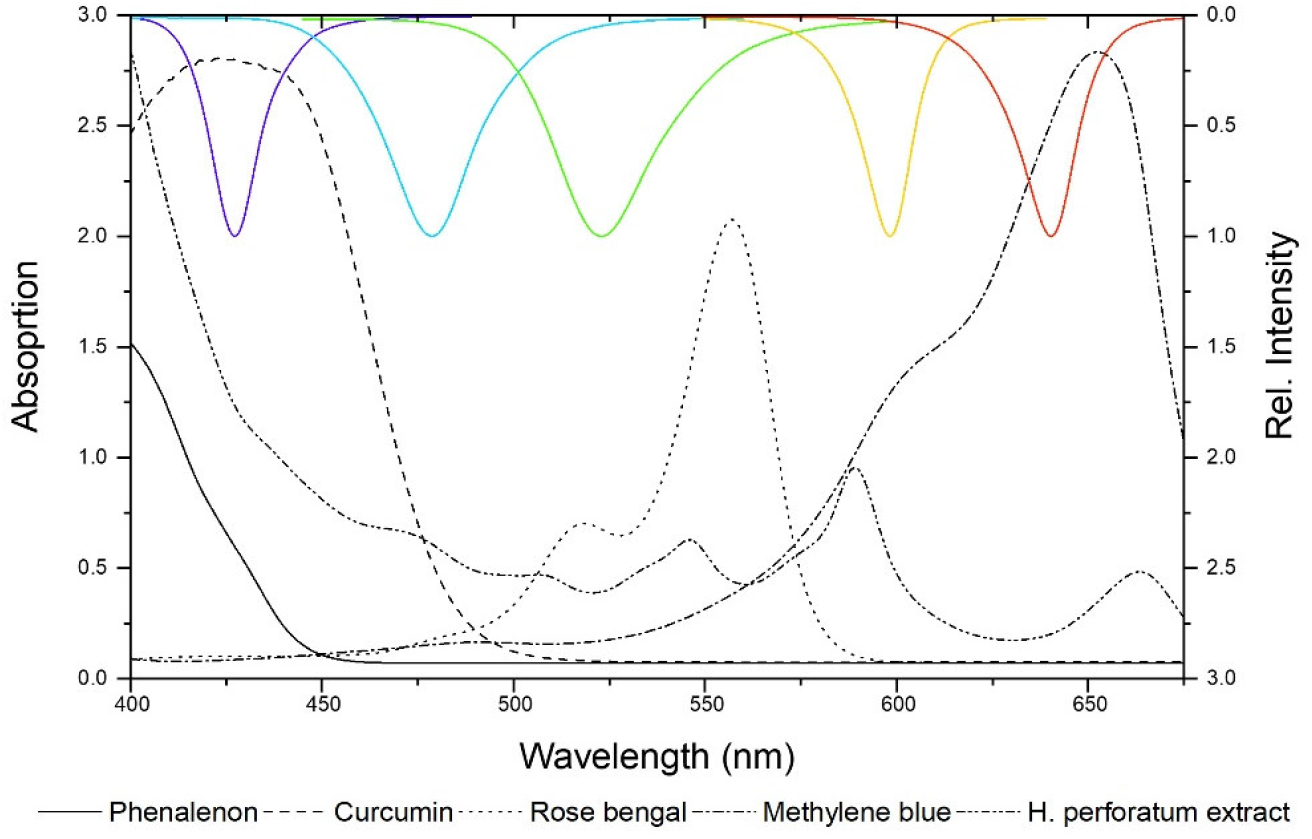
Absorbance spectra of PCs versus emission spectra of the LED modules (λ_irr_ = 428, 478, 523, 598, and 640 nm from left to right).

**Table 2:** Overview of the minimal inhibition concentrations under irradiation (PhotoMIC) of the investigated positive controls regarding the three tested MOs. In the table, for each PS are given the PhotoMIC value with the utilized preincubation time (PI) and light dose (*H*). The last column contains MICs of standard AB without light-irradiation.

|  | 428 nm | 478 nm | 523 nm | 598 nm | 640 nm | Dark |
| --- | --- | --- | --- | --- | --- | --- |
| *C. albicans* (yeast) | Curc<br>4 $\mu\text{g/mL}$<br>(10.9 $\mu\text{M}$)<br>60 min<br>30 J/cm<sup>2</sup> | Curc<br>30 $\mu\text{g/mL}$<br>(81.5 $\mu\text{M}$)<br>10 min<br>9.3 J/cm<sup>2</sup> | HP<br>50 $\mu\text{g/mL}$<br>10 min<br>30 J/cm<sup>2</sup> | HP<br>200 $\mu\text{g/mL}$<br>10 min<br>9.3 J/cm<sup>2</sup> | MB<br>2.5 $\mu\text{g/mL}$<br>(7.8 $\mu\text{M}$)<br>60 min<br>30 J/cm<sup>2</sup> | AMP<br>0.2 $\mu\text{g/mL}$<br>(0.2 $\mu\text{M}$) |
| *E. coli* (gram negative) | Curc<br>40 $\mu\text{g/mL}$<br>(108.6 $\mu\text{M}$)<br>10 min<br>30 J/cm<sup>2</sup> | PN*<br>75 $\mu\text{g/mL}$<br>(416.2 $\mu\text{M}$)<br>10 min<br>9.3 J/cm<sup>2</sup> | RB*<br>150 $\mu\text{g/mL}$<br>(154.1 $\mu\text{M}$)<br>10 min<br>30 J/cm<sup>2</sup> | n.s. | n.s. | CAP<br>2 $\mu\text{g/mL}$<br>(6.2 $\mu\text{M}$) |
| *S. aureus* (gram positive) | Curc<br>4 $\mu\text{g/mL}$<br>(10.9 $\mu\text{M}$)<br>10 min<br>9.3 J/cm<sup>2</sup> | PN<br>25 $\mu\text{g/mL}$<br>(138.7 $\mu\text{M}$)<br>10 min<br>9.3 J/cm<sup>2</sup> | RB*<br>4 $\mu\text{g/mL}$<br>(4.1 $\mu\text{M}$)<br>10 min<br>30 J/cm<sup>2</sup> | HP<br>150 $\mu\text{g/mL}$<br>10 min<br>9.3 J/cm<sup>2</sup> | n.d. | ERY<br>1 $\mu\text{g/mL}$<br>(1.3 $\mu\text{M}$) |

Curc = Curcumin, PN = Phenalenone, RB = Rose bengal, HP = *Hypericum perforatum* extract; \* other PS worked as well. N.d. = Not detected.; n.s. = not selective; CAP = Chloramphenicol; AMP = Amphotericin B; ERY = Erythromycin. For a full discussion see SI Chapter 1.

### 3.4 Mycochemical Analysis of Selected Cortinarius species

Based on their colorful appearance, the fruiting bodies of six different *Cortinarius* species (i.e., *Cortinarius rufo-olivaceus, C. tophaceus, C. traganus, C. trivialis, C. venetus*, and *C. xanthophyllus*) were selected to evaluate our photo-antimicrobial assay (See Table S1 for collection information). In a first step, the dried material was extracted and the obtained extracts (see Table 3) were analyzed spectroscopically (UV-Vis, Figure S7) as well as chromatographically (i.e., HPLC combined with several hyphenated detectors (i.e., DAD, FLD, ELSD, MS see Figure S8-S10)). The results showed that the extract of *C. xanthophyllus* is not only the most complex but also the most intensely colored one (Figure S7 and S11). In detail, five intense peaks were detected at λ = 254 nm (Table S3). The absorption maxima of the two major peaks (t_r, Peak_ 4 = 28.3 and t_r, Peak_ 5 = 34.2 min) were detected and equaled λ_max, Peak_ 4 = 436 nm and λ_max, Peak_ 5 = 525 nm (See Figure S12 UV-Vis spectra).

**Table 3:** Initial weight of biomaterial of *Cortinarius* species and yield of extracts.

|  | Biomaterial<br>[mg] | Yield of extract<br>[mg, (%)] | Visual appearance<br>extract |
| --- | --- | --- | --- |
| *C. rufo-olivaceus* | 1784.9 | 91.0 (5.1) | Dark red, dull |
| *C. venetus* | 1944.0 | 26.7 (1.4) | Light yellow, greasy, |
| *C. tophaceus* | 1643.7 | 14.6 (0.9) | Light yellow, muddy |
| *C. traganus* | 1709.7 | 18.2 (1.1) | Dark yellow, muddy |
| *C. trivialis* | 1958.1 | 22.0 (1.1) | Dark yellow, greasy |
| *C. xanthophyllus* | 1051.8 | 26.6 (2.5) | Purple, earthy, powder |

The mass spectrometric analysis revealed a mass of m/z = 283 [M-H]^−^ for Peak 4 and the chemical formula C_16_H_12_O_5_. Taking the characteristic fluorescence properties of Peak 4 (Figure S8A) and the TLC work of Hofbauer (Hofbauer, 1983) into account, this peak was annotated as parietin and confirmed by comparison with an authentic sample (Figure S13). Also, Peak 2, 3, and 5 were characterized by anthraquinone-like absorption spectra (Figure S12). The red shift of the absorption maxima (Δλ= 57-87 nm, as compared to Peak 4 (parietin)) indicated an extended chromophore and thus hinted towards dimeric AQ-like structures. While Peak 2 (tr = 25.4 min) and Peak 3 (tr = 26.9 min) were also detected in the extract of *C. rufo-olivaceus*, they were putatively assigned as rufoolivacin A & C (Gill and Steglich, 1987; Zhang et al., 2009; Gao et al., 2010), which was in accordance with their mass peak of m/z = 557.2 [M+H]^+^ (Table S3). Peak 5 (m/z = 556.2 [M+H]^+^) was not assigned yet, but might be an oxidated derivative of phlegmacin (MW = 576.6 g/mol), which was described in *C. xanthophyllus* (Hofbauer, 1983). Plenty of dimeric anthraquinones are known from related Cortinariaceae (Gill and Steglich, 1987; Elsworth et al., 1999; Zhang et al., 2009; Gao et al., 2010) and thus seems to be a reasonable putative annotation. Further discussion of the metabolic profiles can be found in the supplementary part (Chapter 2.2.3).

### 3.5 Singlet-Oxygen Detection assay (DMA-Assay)

The obtained extracts were submitted to the recently developed singlet oxygen high-throughput assay (DMA-assay, (Siewert et al., 2019)). Out of the six investigated extracts, two, namely *C. xanthophyllus* and *C. rufo-olivaceus*, showed the ability to produce ^1^O2 after being irradiated with blue light (Table 4). *C. xanthophyllus* was the most active extract: Irradiated at λ = 468 ± 27 nm (24.8 J/cm^2^), the extract produced 187% singlet oxygen as compared to the well-known photosensitizer phenalen-1-one (Schmidt et al., 1994; Espinoza et al., 2016). Hence, this extract originating from natural sources showed promising photosensitizing activity as promising as those of synthetic compounds, such as phenalene-1-one.

**Table 4:** Results of the DMA-assay (blue light irradiation relative to phenalene-1-one).

|  | Singlet Oxygen Production [%] | Standard deviation [%] |
| --- | --- | --- |
| *C. rufo-olivaceus* | 49.6 | 2.55 |
| *C. tophaceus* | 3.1 | 1.7 |
| *C. traganus* | 0.2 | 1.1 |
| *C. trivialis* | 4.2 | 2.0 |
| *C. venetus* | 4.4 | 0.2 |
| *C. xanthophyllus* | 187.1 | 2.1 |

### 3.6 *Cortinarius xanthophyllus* contains photoantimicrobials active against *Staphylococcus aureus* and *Candida albicans*

Submitting all six extracts to the (photo)antimicrobial assay revealed that all extracts are inactive (c > 50 μg/mL) under the exclusion of light (Figure 6-8). Under light-irradiation, however, the activity of the purple extract of *C. xanthophyllus* was significantly enhanced: The growth of the gram-positive bacterium *S. aureus* (Figure 8) was completely inhibited with an extract concentration as low as c = 7.5 μg/mL and a light dose of H = 30 J/cm^2^ (λ = 478 nm). This also holds true for the photoactivity against the yeast *C. albicans*, where an extract concentration of c = 75 μg/mL (H = 30 J/cm^2^) led to complete growth inhibition (Figure 6). Against the gram-negative *E. coli*, however, *C. xanthophyllus* was inactive in the dark and under irradiation (Figure 7).

**Figure 6:**
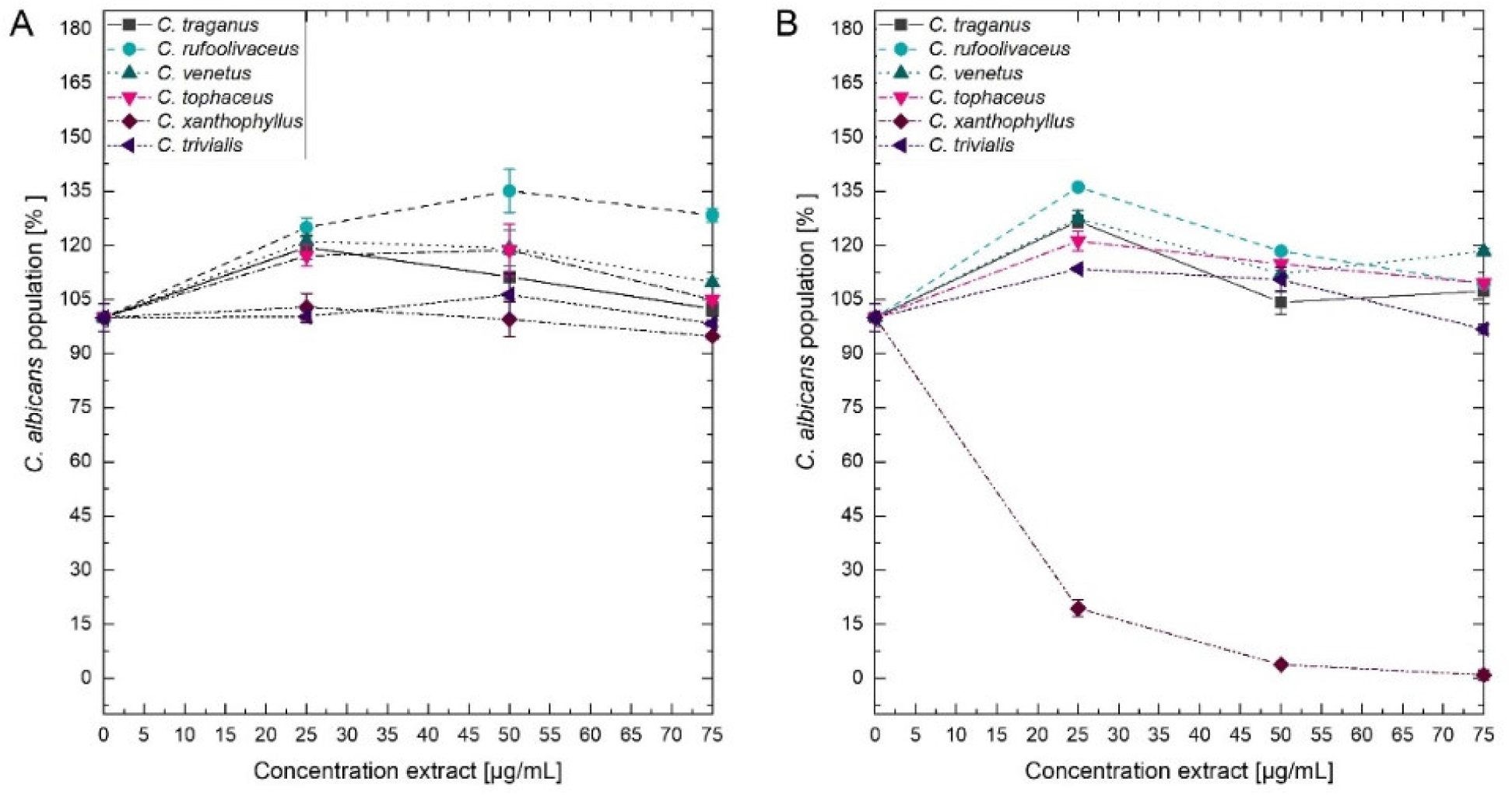
Dose-response curves of *C. albicans* treated with *Cortinarius* extracts. On the left side (A) dark controls are shown. The right graph (B) represents the irradiated experiments (λ = 470 nm, H = 30 J/cm^2^, PI = 60 min).

**Figure 7:**
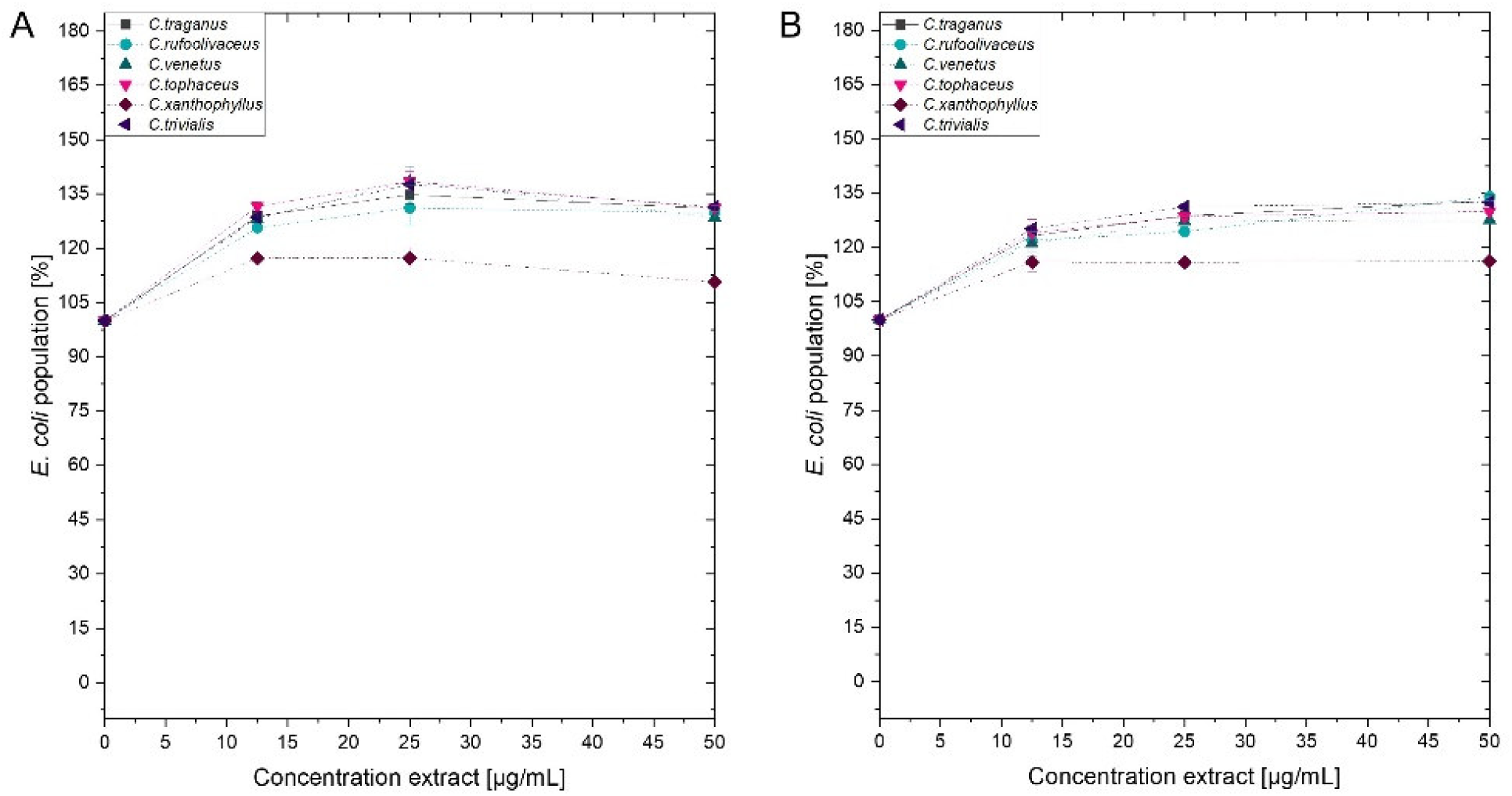
Dose-response curves of *E. coli* treated with *Cortinarius* extracts. On the left side (A) dark controls are shown. The right graph (B) represents the irradiated experiments (λ = 470 nm, 30 J/cm^2^, PI = 60 min).

**Figure 8:**
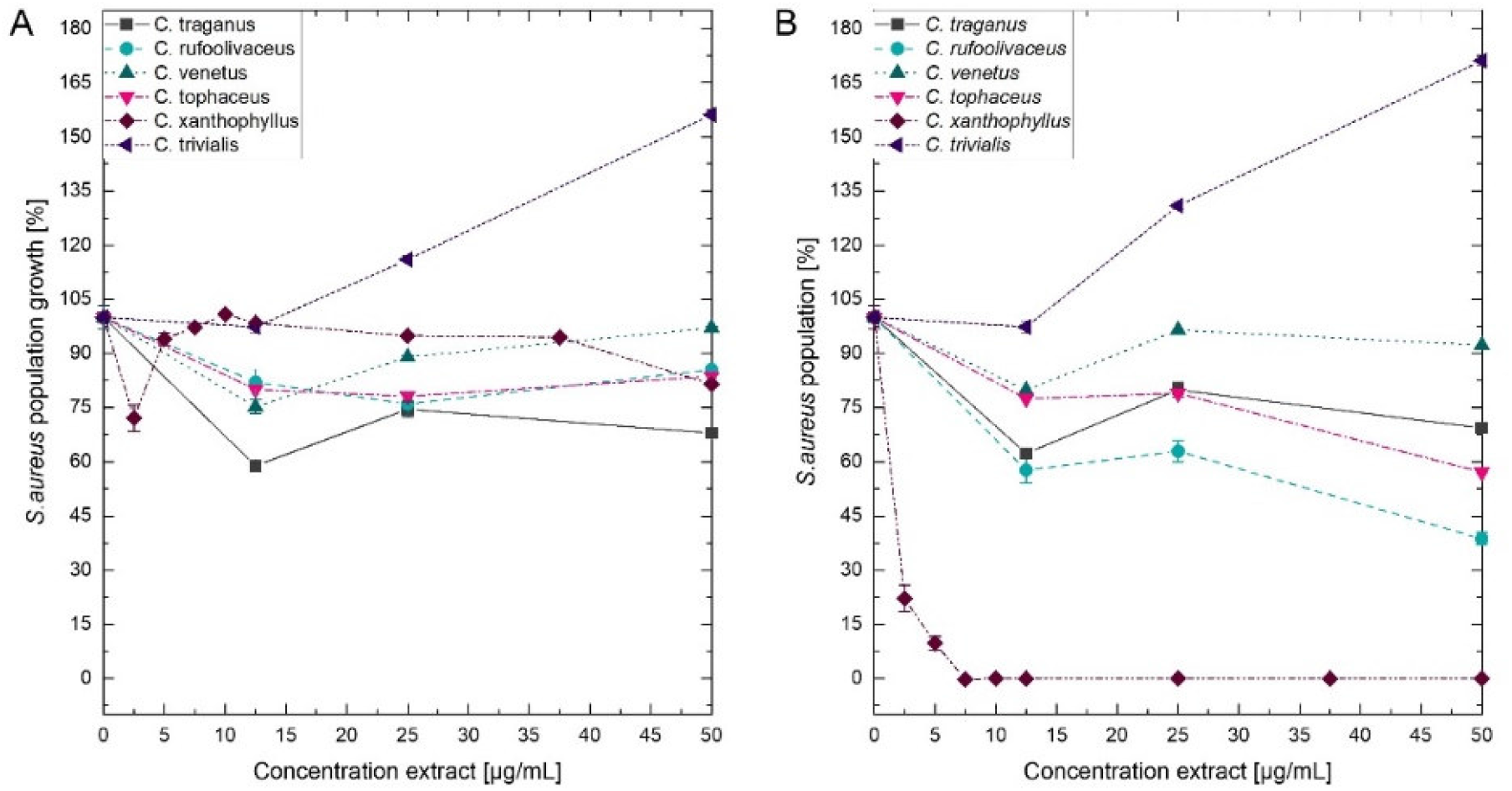
Dose-response curves of *S. aureus* treated with *Cortinarius* extracts. A) Growth inhibition under dark conditions. B) Growth inhibition under irradiation conditions (λ = 470 nm, H= 30 J/cm^2^, PI = 60 min).

A weak enhancement of the growth inhibition effect was also seen for the *C. rufo-olivaceus* extract against *S. aureus* (IC_50_ ~ 50 μg/mL). Nevertheless, this extract did not affect *C. albicans* under the tested conditions.

## 4 Discussion

The hypothesis we wanted to test in course of this study was that light is a neglected co-factor in antimicrobial screening assays. Therefore, a convenient HTS-screening assay based on the EUCAST protocol was established and validated. In a first step, an innovative LED-panel was required achieving a homogenous irradiation of a 96-well plate. Finally, the hypothesis was be tested with a set of six different Cortinarius species.

### 4.1 The SciLED irradiation system and its improved uniform irradiance distribution

To achieve the first objective – the homogenous irradiation of a 96-well plate – the distance and number of the LEDs was optimized by simulations until the coefficient of variation *cv* was estimated to be less than one percent. Irradiance measurements and chemical actinometer measurements (Figure 3) confirmed the homogeneous irradiance. Nevertheless, the actual coefficient of variation from irradiance measurements was in the range between seven to nine percent. Deviations between the simulation and the measurement can be attributed to differences between the modeled and the actual radiant intensity distribution of the LED, variations in the radiometric power of individual LEDs, and measurement uncertainties.

Although the real uniformity (*cv* = 8%) of the irradiation system with non-equidistant LED arrangement presented in this work was less than the expected uniformity from the simulations (*cv_sim_* = 0.08%), the achieved homogeneity over the whole sample area was still significantly better compared to equidistant LED positioning. An optical simulation of a 6 × 4 LED array with an equidistant arrangement (Δ*x* = 35 mm, Δ*y* = 35 mm) resulted in a less homogeneous irradiance distribution (*cv_sim_* = 5%) compared to the existing non-equidistant arrangement (*cv_sim_* = 0.08%). To understand the positive effects of a non-equidistant positioning, the non-uniform radiant intensity distribution of LEDs must be taken into account. Considering just one LED, the resulting irradiance distribution on the irradiated plane is decreasing nonlinear with an increased distance from the center. Placing two LEDs with a certain distance *d* next to each other, parts of the irradiance will overlap. Due to the superposition principle, the resulting irradiance distribution is the sum of every single one (Figure 9). Depending on the distance, the irradiance in the intermediate area between the two LEDs is enhanced, reduced, or constant. This dependence of the irradiance distribution from the LED distance is shown in Figure 9 for three different distances *d*. Using an array of *n* × *m* LEDs with *d* > 2 and *m* ≥ 1, the overlapping effect is amplified. To achieve a uniform distribution, the right LED arrangement is crucial. The LED distances for a uniform irradiance depends on the number of LEDs, the viewing angle *θ*_1/2_ and the working distance *z*. As a rule of thumb, for LEDs with a wide viewing angle, the distances of the inner LEDs should be wider than the distance of the outer ones.

**Figure 9.**
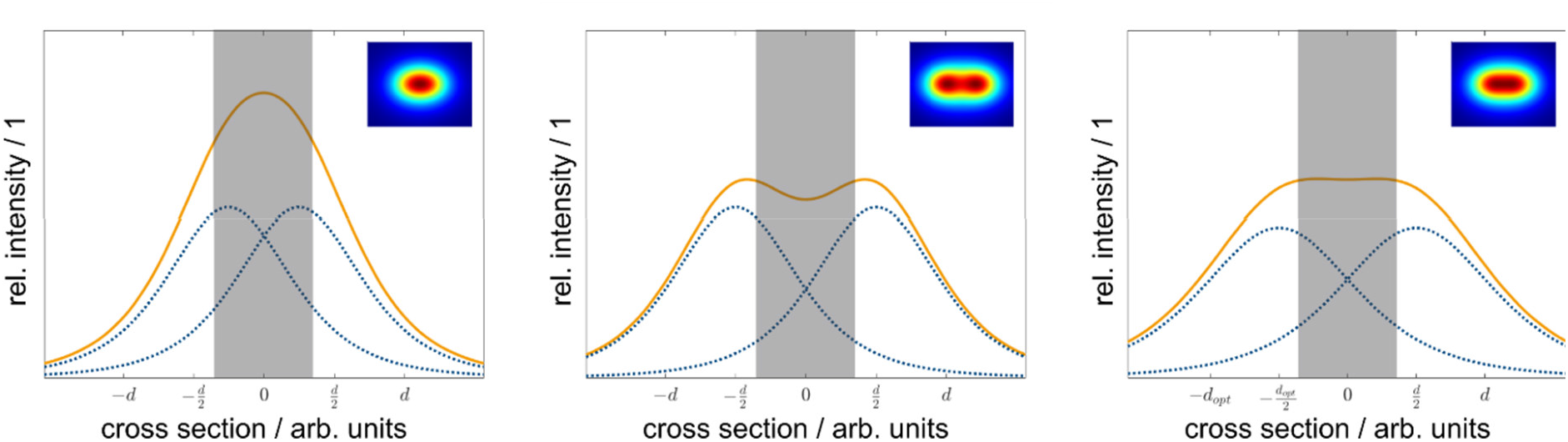
Side-view of the irradiance distribution depending on LED distance. Depending on the LED distance ***d***, the resulting irradiance from the two overlapping LED irradiance distributions at the sample plane differs. Individual LED irradiance distributions are represented as dotted lines (blue), and the resulting distributions are shown as solid lines (orange) The grey bar indicates the max. width of the sample plate. If the LED distance ***d*** is too low, the overlap in the center is high, and the resulting irradiance is enhanced (A). For a too large LED distance ***d***, the overlap is small, and the resulting irradiance is reduced (B). Using the optimal LED distance ***d_opt_***, each LED contributes the right amount, and the resulting irradiance in the center is constant (C).

However, a uniform irradiance comes with a price. Due to the positioning scheme, the average irradiance decreased by ten percent compared to the equidistant arrangement. Simulations of different equidistant LED positions have shown that the resulting average irradiance increased by reducing the spacing between the individual LED, yet the uniformity decreased. Depending on the purpose of the irradiation system, a tradeoff between irradiance and homogeneity is necessary. For this work, a uniform distribution was essential to accomplish comparable irradiation conditions within the 96-well plate.

Optical characterization measurements shown in Table 1 indicate that the actual peak wavelengths are within the specifications from the datasheets for all but the red LEDs. With a nominal wavelength range from λ = 624 nm to λ = 634 nm given by the manufacturer and an actual peak wavelength of λ = 640 nm a divergence was observed. This deviation may result from a different characterization method in the datasheet. The LED manufacturer refers to a dominant wavelength, which takes the relative spectral sensitivity of human visual perception of brightness (luminosity function) into account (Lumileds Holding B.V. 2019). The peak wavelength in this work refers to an absolute spectral measurement without considering the luminosity function of the human eye. Irradiation measurements at the sample distance revealed a variation in the achieved intensities from a maximum irradiance of *E_m_* = 13 ± 1.0 mW/cm^2^ for violet LEDs and a minimum irradiance of *E_m_* = 1.1 ± 0.08 mW/cm^2^ for amber LEDs. These fluctuations can be explained by different designs and composition of each single-color LED. To achieve different emission wavelengths, different semiconductor combinations with different dopings are used in addition to various packaging layouts (Schubert, 2006). These intrinsic variations result in different irradiances.

### 4.2 Establishment and Validation of a Screening Photoantimicrobial Assay

The EUCAST microdilution assay – being launched to allow a better inter-laboratory reproducibility – inspired the established photoantimicrobial HTS. While antimicrobial assays heavily depend on the testing conditions, one aim of EUCAST is to boost the development of new antimicrobials by the enablement of inter-laboratory comparisons. Specifically, photoantimicrobials are part of a promising treatment alternative, the so-called Photoantimicrobial chemotherapy (PACT) or antimicrobial photodynamic inhibition (aPDI) (Wainwright, 2009). While the therapy depends on a completely different mechanism (i.e., ROS production due to the interplay of light and a photosensitizer) compared to well-established antibiotics, it is active against multi-resistant pathogens and the risk of resistance-development is relatively low (Maisch, 2015). Nevertheless, despite these attractive aspects, wide acceptance of PACT is lacking. One limitation next to others (Wainwright, 2009) is the limited throughput of established assays: Often, one PS-candidate and one concentration are irradiated by the time, leading to exorbitantly time consuming experiments. On the other hand, testing multiple parameters (concentrations, microorganism, or drug-candidates) on one plate lacked comparability due to an uneven light distribution (Ogonowska et al., 2019). Thus, the limited throughput impeded the study of extensive libraries of PS-candidate and hence the classical lead-to-hit approach of medicinal chemistry.

The EUCAST protocol tests an antibiotic (AB) usually with ten concentrations and defines the MIC by the value which is the lowest concentration inhibiting the growth completely as determined by the lack of turbidity. To test photoantimicrobials, a blank measurement of each concentration is necessary. The test solutions are colored, and the blank subtraction is necessary to avoid false-negative read-outs during the OD measurement. Furthermore, a triplicate of each concentration is needed account for the biological variability. This led us to the, in Figure S8 displayed, pipetting scheme, which allows testing the effect of two PSs against one microorganism. While for classic EUCAST susceptibility assays only these variables (i.e., tested microorganism and concentration range of the AB) are crucial, the number of variables exceeds in the photoantimicrobial assay: In addition to the preincubation time, irradiation time, and light doses, as well as light power and the irradiation wavelength itself are of interest.

The established scheme (Figure 2) and the workability were tested with the known PSs curcumin, rose bengal (RB), methylene blue (MB), phenalenone (PN), and a *Hypericum* extract (HP). The irradiation wavelength changed according to the absorbance pattern of the PS (Figure 5). The light dose was set to *H* = 30 J/cm^2^, which equated to the utilized dose from several published studies (Cieplik et al., 2016; de Annunzio et al., 2018; Merigo et al., 2019) and furthermore was shown to be non-toxic against the tested microorganisms alone (Figure 4). While this is per se not as important as for photocytotoxicity (Wainwright, 2009) studies, we choose this dose to truly see the light-effect of the PSs. PhotoMIC values were generated for this variety of PSs against the pathogenic microorganisms *S. aureus, C. albicans*, and *E. coli* utilizing the LED modules with an irradiation wavelength of λ_irr_ = 428, 478, 523, 598, and 640 nm. To the best knowledge of the authors, Table 2 represents the first comprehensive overview of the PhotoMIC values of common PSs. Although literature values cannot be easily compared due to the discussed, yet not standardized parameters (Haukvik et al., 2009), our obtained values fit the reported ones (See Supplementary Part, Chapter 1.2 for the full discussion). An international agreement on standard values for the additional irradiation parameters would be helpful in the process of hit-lead optimization.

The gram-negative bacteria *E. coli* was resistant against the tested lipophilic and neutral PSs especially during the irradiation with yellow and red light (Table 2). This is well-known (see discussion SI chapter 4.1) and reasoned by their negatively charged membrane (Minnock et al., 2000; Bresolí-Obach et al., 2018; Galstyan et al., 2018).

To allow a screening of biological sources such as plant extracts or fungal extracts in the frame of bio-activity guided isolation, the pipetting scheme displayed in Figure 2 was established. Due to the even light distribution, up to seven extracts à three concentrations can be screened against one pathogenic microorganism in biological triplicates. By a slight modification of the testing logic on the other hand, a fast determination of PhotoMIC against a broader variety of microorganisms in analogy to the EUCAST scheme is possible (i.e., up to seven microorganisms against one PSs, no triplicates).

### 4.3 Utilization of the Screening Assay Yielded a Promising Hit

As a sample set, extracts of six different *Cortinarius* species (Table 3 and Table S1) were investigated. The results of the antimicrobial assay (dark conditions) were in line with the results of Tiralongo and colleagues (Beattie et al., 2010). They investigated 117 different Australian *Cortinarius* species and could show that two-third of the species held an IC_50_ between c = 20 μg/mL and c = 200 μg/mL against the gram-positive bacterium *S. aureus*. In the present study, we determined MICs (instead of IC_50_), and were, under light exclusion, not able to see full inhibition of microbial growth with extract concentrations up to c = 75 μg/mL.

As shown in Figure 6, the addition of blue light exhilarated the antimicrobial activity of the intensely colored *Cortinarius* extract by more than tenfold: The extract of *C. xanthophyllus* was characterized by a PhotoMic of c = 7.5 μg/mL and thus was even more effective than the established PS phenalenone (MIC = 25 μg/mL). In addition, the extract showed promising activity against *C. albicans* (Figure 6). Interestingly, this activity seems to be uptake depended, as a preincubation time of only ten minutes (instead of PI = 60 min) showed no effect (Figure S14). Mycochemical analysis of the extract implicated three potential photoactive compounds (Figure S11). These pigments were tentatively annotated as rufoolivacin, parietin, and as an anhydro-phlegmacin-like compound (Table S3). Analysis of the HPLC-DAD chromatogram recorded at λ = 478 nm indicated that Peak 4 and Peak 5 absorb most of the incoming light, and thus might be responsible for the observed photoantimicrobial action. Parietin, usually isolated from the lichen *Xanthoria parietina* is known for its photoactive properties against cancer cells (Mugas et al., 2021) and against *S. aureus* (Comini et al., 2017). The chemical structure of Peak 5 is not assured yet and hampered by the limited availability of fungal material due to the rare occurrence of *C. xanthophyllus*. This Mediterranean species is listed on the red-list and thus endangered (Tingstad et al., 2017). Nevertheless, applying modern phytochemical techniques (e.g., LC-SPE-NMR, FBMN-assisted isolation) might help to reveal its chemical space and is part of future work.

## 5 Conclusion

The development of an uniform emitting LED-panel was presented, allowing a homogeneous irradiation of a complete 96-well plate. As consequence, a convenient HTS-assay to determine photo-activated minimal inhibitory concentrations (PhotoMIC) of pure compounds and extracts was established based on the EUCAST guideline. The light tolerance of the utilized model organisms (i.e., *C. albicans, E. coli*, and *S. aureus*) was tested and revealed that all microorganisms can cope with a light dose of *H* = 9.3 J/cm^2^ or even *H* = 30 J/cm^2^ of every tested wavelength (i.e., up to 9.3 J/cm^2^ for λ = 598, up to 30 J/cm^2^ for λ = 428, 478, 528, 640 nm). Standard photosensitizers were used to validate the assay and yielded the first comprehensive table accumulating a broad array of PhotoMic values under different irradiation conditions and against different pathogenic MOs. Lastly, submitting a test sample set of fungal extracts generated from the colored fruiting bodies of *Cortinarius rufo-olivaceus, C. tophaceus*, *C. traganus*, *C. trivialis*, *C. venetus*, and *C. xanthophyllus* showed that light is indeed one co-factor amplifying a moderate antimicrobial action of natural products. The most intensely colored extract, i.e., the one of *C. xanthophyllus*, showed the most promising activity of PhotoMIC = 7.5 μg/mL against *S. aureus*. The extract was also photoactive against *C. albicans*. Mycochemical analysis identified two peaks putatively responsible for the effect. One of them being the well-known natural PS parietin from the lichen *Xanthoria parientina* and the other one being a photochemically unexplored dimeric AQ.

## 7 Author Contributions

J.F. performed the antimicrobial assays and majority of the mycochemical analysis. F.H. performed the DMA-assay. H.S. and R.S. designed the irradiation device. H.S. performed the instrumental characterization. D.D. and D.J.A. performed pre-test of the AntiMic assay. P.V. contributed to the conception of the AntiMic assay. U.P. provided the biomaterial and phylogenetic input. B.S. designed the research, analyzed the mycochemical part, and wrote the manuscript with contributions of H.S. and J.F. All authors contributed to the final version of the manuscript.

## 9 Acknowledgments

H. Stuppner is kindly acknowledged for his support and input. Sarah Flatscher is acknowledged for her skillful help characterizing the LEDs.

## 10 Supplementary Material

See attached document.

