## Supplementary Information for "A new High-Throughput-Screening-assay for Photoantimicrobials Based on EUCAST Revealed Photoantimicrobials in Cortinariaceae"

*Supplementary Material*

### 1 Photoantimicrobial Assays

#### 1.1 Pipetting Scheme for photoantimicrobials

In Figure S1 a pipetting scheme for photoantimicrobials is presented.

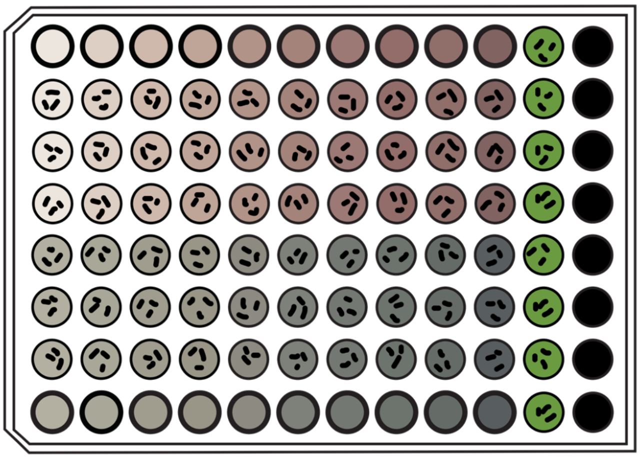

**Figure S1** Pipetting scheme for two antimicrobials (Compound A = brown and B = grey) tested against one microorganism in ten different, from left to right rising, concentrations. Bold strokes imply the blank of each tested concentration. GC = Growth control = green vials, SC = Sterility control = full black vials.

#### 1.2 (Photo)antimicrobial action of positive controls

##### 1.2.1 Curcumin

Our experimental photoMIC for curcumin (c = 4 µg/mL, 10.9 µM, *H* = 30 J/cm², λ = 428 ± 15 nm, 60 min PI, Figure S2) against *Candida albicans* concurred with the findings of Dovigo and colleagues(Carmello et al., 2015;2017), who reported the minimum fungicidal concentration (MFC) for curcumin (c = 20 µg/mL, 54.3 µM). They utilized a similar light dose (*H* = 37.5 J/cm$^2$) and a comparable light source (λ = 455 -15/+5 nm) but a three times shorter PI (t = 20 min). The potential influence of PI for PACT against *C. albicans* can also be seen in the results of *C. xanthophyllus* against *C. albicans* (Figure S14). Although MFCs are higher than MICs, the results are moderately comparable. In line with the data from Dovigo (2015), irradiation with λ = 478 ± 28 nm light and a fluence of 9.3 J/cm$^2$ increases the MIC to c = 30 µg/mL, 81.4 µM (Table 2, Figure S4).

The reported photoantimicrobial action of curcumin against the gram-negative bacteria *Escherichia coli* are quite controversial: (Bhavya and Umesh Hebbar, 2019) found a 5.94 log CFU/ml reduction (λ = 462 nm, *H* = 13 J/cm$^2$, growth condition = 75 mm petri dish) with a concentration of c = 7.37 µg/mL, (c = 20.0 µM). In their experiment it was reported that a PI up to t = 60 min did not show any significant difference on the results.

De Oliveira et al. report a reduction up to 3 log CFU/mL at a concentration of c = 5 µg/mL (13.57 µM) and a smaller reduction of 1.8 log CFU/mL with a concentration of c = 10 µg/mL (27.15 µM) (Medium = citrate buffer (c = 100 µM), growth conditions = 12-well flat bottom polystyrene plate, light source = UV-A lamps, λ = 320–400 nm, PI = 5 min) (de Oliveira et al., 2018). In another report, reductions of just 1.29 and 2.65 log CFU/ml were found when *E. coli* was treated with curcumin (c = 27.63 µg/mL, 75 µM) under comparatively strong irradiation conditions (λ = 470 nm, $H$ = 139 and 278 J/cm$^2$, respectively, PI = 10 min) (Penha et al., 2017). Our experimental PhotoMIC ($H$ = 30 J/cm², λ = 428 ± 15 nm, Figure S2) for curcumin against *E. coli* equalled c = 40 µg/mL (c = 108.6 µM).
Reasons for these controversial results might be the different irradiation setups, the preincubation time, the method (i.e., microdilution assay *vs* colony forming assay) itself, or the different susceptibilities of different strains.

Against *S. aureus* a MIC of c = 4 µg/mL (10.86 µM) curcumin (λ = 430 nm, $H$ = 9.3 J/cm², PI = 10 min) was found (Table 2, Figure S2). With a concentration of c = 2 µg/mL (5.43 µM) a 50% inhibition of growth was shown. (Jiang et al., 2014) reported a 2 log CFU/ml reduction at c = 2.5 µM (0.92 µg/mL) with a smaller light dose and a longer PI (λ = 470 nm, H = 3 J/cm$^2$, PI = 60 min).

##### 1.2.2 Phenalenone

As showed by others, we found no photoactivity of phenalenone against *C. albicans* (c = 1.13 µg/mL – 90.10 µg/mL (6.25 – 500 µM), λ = 380 - 500 nm, H = 12 J/cm$^2$, PI = 240 min) (Bauer, 2016). Against *S. aureus* and *E. coli* we found PhotoMICs of c = 25 µg/mL (138.7 µM) and c = 75 µg/mL (416.2 µM), respectively (Figure S2). Phenalenone derivatives are known to be promising natural PSs (Flors and Nonell, 2006). The backbone structure itself, showed no photoantimicrobial activity against *S. aureus, E. faecalis,* and *E. coli* with small light doses and low concentrations (c = 1.80 µg/mL, 10.0 µM), λ = 420 ± 10 nm, $H$ = 1.2 J/cm$^2$, PI = 10 min) (Bresolí-Obach et al., 2018) or was not considered itself (Muehler et al., 2017) (Tabenski et al., 2016). Derivatives of PN, especially those with a positive charge showed promising photoantimicrobial activities (Bresolí-Obach et al., 2018).

##### 1.2.3 Rose bengal

With a concentration of c = 48.69 µg/mL (50.0 µM) rose bengal Rossoni et al. showed a 7.73 log CFU/ml reduction for *E. coli* using a light dose of $H$ = 94.74 J/cm$^2$ (λ = 460 nm (no deviation reported), PI = 5 min) (Rossoni et al., 2010). Our PhotoMIC is c = 150 µg/mL (154.1 µM) using a third of their light dose ($H$ = 30 J/cm$^2$, PI = 10 min). Nisnevitch et al. did not find a MIC below a concentration of c = 194.74 µg/mL (200.0 µM) for rose bengal against *E.coli* (λ = 400 – 700 nm, $H$ = 4.53 J/cm$^2$, no PI reported) (Nisnevitch et al., 2010).
For different strains of *S. aureus* Ilizirov et al. found PhotoMICs between c = 0.625 – 2.50 µg/mL, (0.642 – 2.568 µM, λ = 400 – 700 nm, $H$ = 2.265 J/cm$^2$, no PI reported) (Ilizirov et al., 2018) for rose bengal. Similar concentrations were determined by Nisnevitch and colleagues, who reported MIC values of c = 2.92 µg/mL (3 µM) with comparable irradiation settings (λ = 400 – 700 nm, $H$ = 4.53 J/cm², no PI reported) (Nisnevitch et al., 2010). Employing our PhotoMIC assay we determined a MIC of c = 4 µg/mL (c = 4.1 µM) using a higher light dose (λ = 523 ± 33 nm, $H$ = 30 J/cm$^2$, PI = 10 min, Figure S4Figure S5).

##### *1.2.4 Hypericum perforatum* Extract

With the easily available ethanolic *Hypericum perforatum* extract, we were able to evaluate our assay with the photoactive natural product hypericin. As recently presented by Delcanale and colleagues

(Delcanale et al., 2020), hypericin is highly active against *S. aureus* (reduction of 7 log CFU/mL at a concentration of c = 0.50 µg/mL (1 µM), λ = 515 ± 40 nm, *H* = 9.6 J/cm$^2$, PI = 30 min). This result was confirmed by us (Figure S4 and Figure S5) and in addition we were able to observe the photodynamic inhibition of growth against *C. albicans* at different wavelengths of irradiation (Table 2). Therewith, the observations of Alam et. al could be verified (Alam et al., 2019).

##### *1.2.5 Methylene blue*

For methylene blue (MB) we found a PhotoMIC of c = 2.5 µg/mL (7.8 µM) against *C. albicans* (λ = 640 nm, *H* = 30 J/cm², PI = 60 min, Figure S6). This value fits in between the diverging literature results: Pasyechnikova et al. found a maximal reduction of MO growth with c = 1 µg/mL, 3.1 µM, (λ = 630 nm (no deviation and light dose reported), PI = 30 min) (Pasyechnikova et al., 2009). Hosseine et al. found an inhibitory concentration of c = 0.01 µg/mL (0.03 µM) for their setup (λ = 660 nm, *H* = 19.2 J/cm$^2$, PI = 10 min) (Hosseini et al., 2016). Higher PhotoMICs are reported by Torres-Hurtado et al., who used a concentration of c = 6.40 µg/mL (20.0 µM) of MB (λ = 600 - 650 nm, *H* = 60 J/cm$^2$, PI = 30 min) to achieve a growth inhibition of 99% for *C. albicans* (Torres-Hurtado et al., 2019). Da Collina and colleagues reported 20 µg/mL (62.53 µM) as the optimal concentration for their setup (λ = 640 ± 12 nm, *H* = 4.68 J/cm$^2$) and different PIs (1 - 20 min) did not influence the results (da Collina et al., 2018). Carvalho et al. reported a concentration of c = 50 µg/mL (156.32 µM) as the optimal MB concentration to inhibit 80 - 90% of the *C. albicans* grown after irradiation (λ = 684 nm (no deviation reported), *H* = 28 J/cm$^2$, PI = 5 min) (Carvalho et al., 2009).

Against *E. coli* Peloi et al. found a concentration of c = 11.3 µg/mL (35.2 µM) (λ = 665 -65/+35 nm, *H* = 4 J/cm$^2$, no PI reported) leading to 93.7% inhibition of growth (Peloi et al., 2008). Wainwright et al. induced total cell death with a three-times higher concentration (c = 31.4 µg/mL, 100 µM) of MB (*H* = 6.3 J/cm$^2$, no further details reported) (Wainwright et al., 1997). Our findings of a PhotoMIC for MB against *E. coli* are to be found in between the reported results (Table 2). We decided not to use MB as a PC against *E. coli* because the light enhanced effect was too small (i.e., light toxicity (c = 20 µg/mL, 63.7 µM, λ = 640 ± 18 nm, *H* = 30 J/cm$^2$, PI = 10 min) *vs* dark toxicity (c = 30 µg/mL, 95.5 µM)).

The dark toxicity of MB against *E. coli* was recently discussed, raising the question of resistance development in certain strains (Gunics et al., 2000;Thesnaar et al., 2021). Overlapping light and dark toxicity for MB against several MOs is known, such as *Enterococcus faecalis* and *Enterococcus faecium* (Wainwright et al., 1999) or *S. aureus* (Wainwright et al., 1998) (Wainwright and Crossley, 2002). For antimicrobial activity of MB against *S. aureus* in the dark the MIC was reported with a concentration of c = 16 µg/mL (50.0 µM) (Thesnaar et al., 2021). Future research should investigate if a lower concentration of MB with a higher light dose and an experimental setup with stricter exclusion of surrounding light extends the therapeutical window of MB.

#### 1.3 Dose-response curves of tested MOs against PCs with different wavelengths of light irradiation

##### 1.3.1 Dose-response curves of irradiation with blue-light

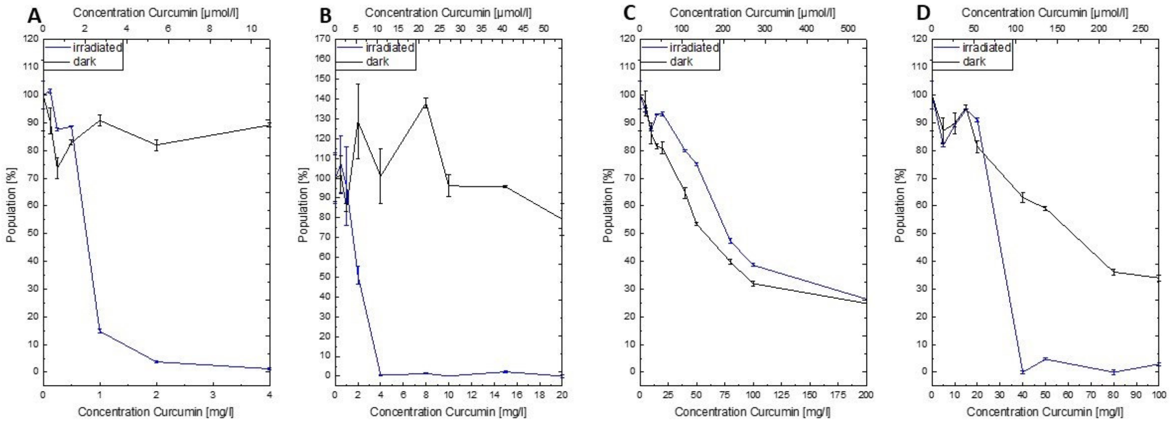


**Figure S2**. (Photo)antimicrobial action of curcumin against A) *C. albicans* (λ = 430 nm, *H* = 30 J/cm$^2$, PI = 60 min)*,* B) *S. aureus* (λ = 430 nm, *H* = 9.3 J/cm$^2$, PI = 10 min), C) *Escherichia coli* with a lower light dose (λ = 430 nm, *H* = 9.3 J/cm$^2$, PI = 10 min,), and D) *E. coli* at a higher light dose (λ = 430 nm, *H* = 30 J/cm$^2$, PI = 10 min).

##### 1.3.2 Dose-response curves of irradiation with purple-light

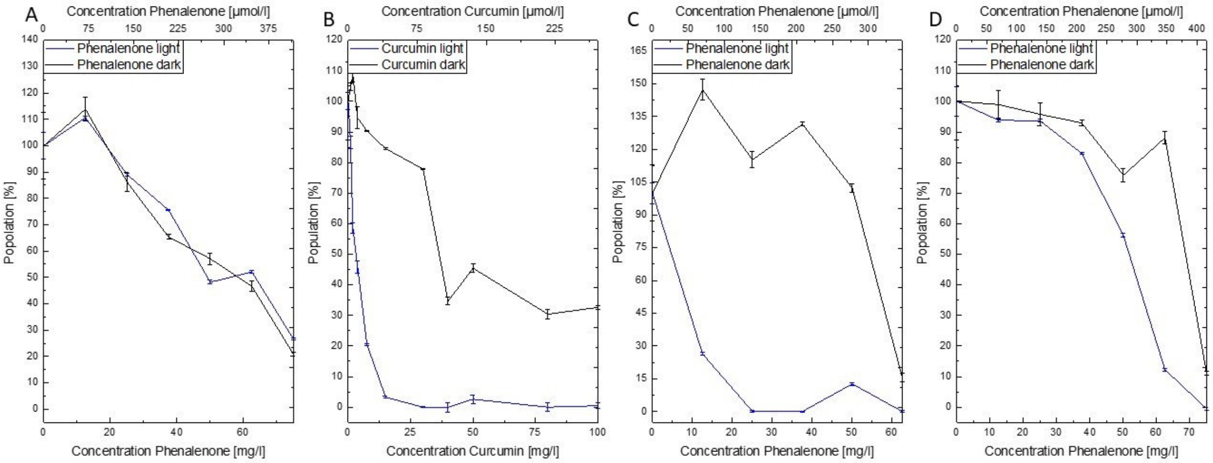


**Figure S 3** (Photo)antimicrobial action of phenalenone (A, C, D) and curcumin (B) used as PC against A) *C. albicans* (λ = 478 nm, *H* = 9.3 J/cm$^2$, PI = 10 min)*,* B) *S. aureus* (λ = 478 nm, *H* = 9.3 J/cm$^2$, PI = 10 min), C) Phenalenone against *S. aureus* (λ = 478 nm, H = 9.3 J/cm$^2$, PI = 10 min), D) Phenalenone against *E. coli* (λ = 478 nm, H = 9.3 J/cm$^2$, PI = 10 min).

##### 1.3.3 Dose-response curves of irradiation with green-light

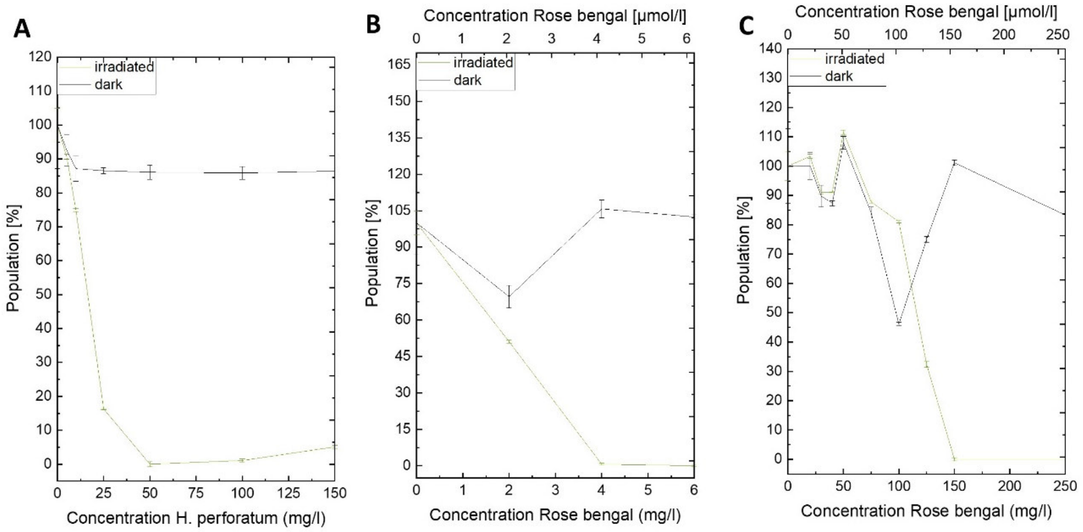


**Figure S4.** Dose-response curves recorded for A) HP against *C. albicans* ($\lambda$ = 523 nm, $H$ = 30 J/cm$^2$, PI = 10 min), B) RB against *E. coli* ($\lambda$ = 523 nm, $H$ = 30 J/cm$^2$, PI = 10 min), C) RB against *S. aureus* ($\lambda$ = 523 nm, $H$ = 30 J/cm$^2$, PI = 10 min).

#### 1.4 Dose-response curves of irradiation with orange-light

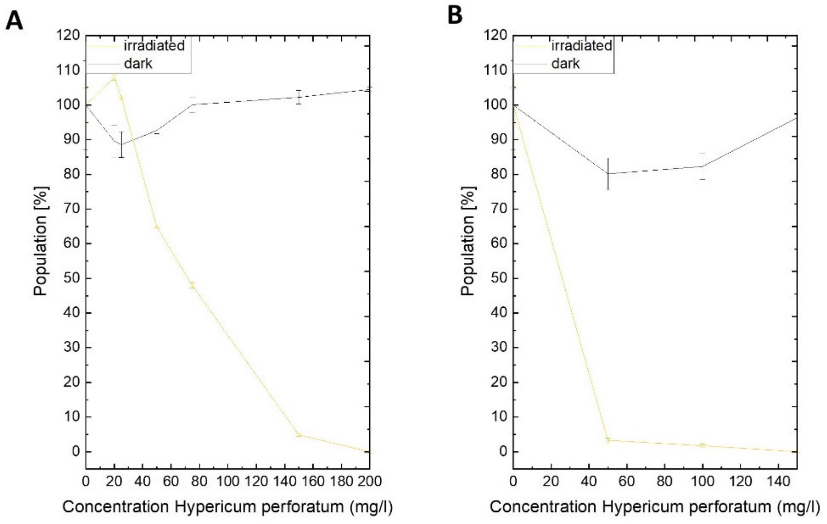


**Figure S5.** Dose-response curve of HP against A) *C. albicans* and B) *S. aureus*. All MOs were preincubated for PI = 10 min and irradiated with orange light ($\lambda$ = 598 nm, $H$ = 9.3 J/cm$^2$).

##### 1.4.1 Dose-response curves of irradiation with red-light

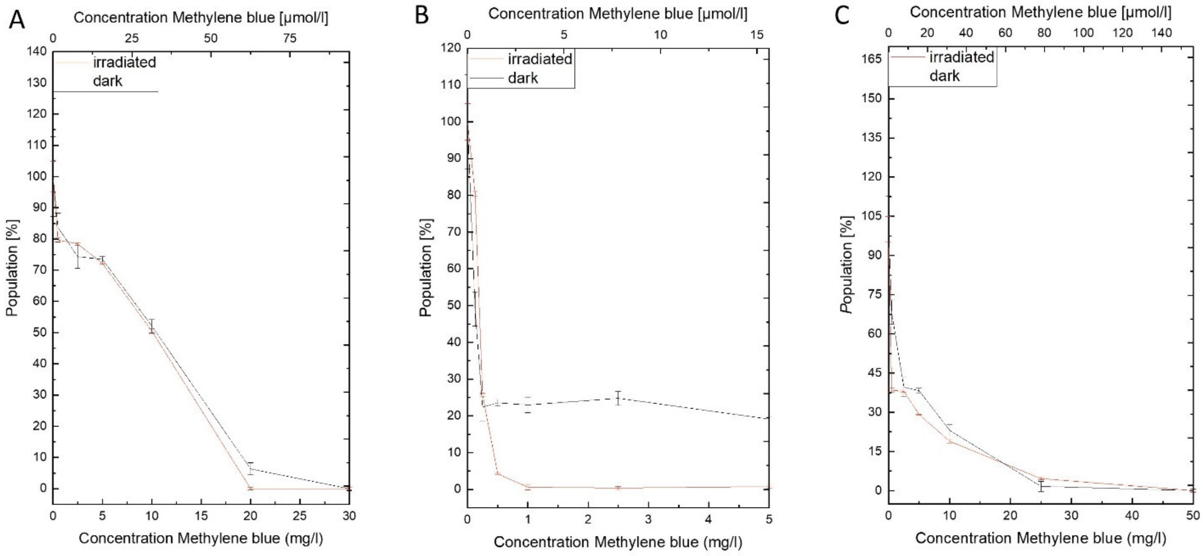


**Figure S6.** Dose-response curve of MB against A) *C. albicans*, B) *E. coli*, and C) *S. aureus*. All MOs were preincubated for PI = 10 min and irradiated with red light ($\lambda$ = 630 nm, $H$ = 30 J/cm$^2$).

### 2 Mycochemical Part

#### 2.1 Fungal material

**Table S1.** *Cortinarius* collections used in this study with respective voucher numbers and collection data.

| | Voucher | leg. et det. | Origin |
|---|---|---|---|
| ***C. traganus*** | IBF20170556 | 14.10.2017 | Pian di Carniglia, Bedonia |
| ***C. rufoolivaceus*** | IBF20190113 | 20.10.2019 | Oasi Ghirardi, Bedonia, Italy |
| ***C. venetus var. montanus*** | IBF20180223 | 15.10.2018 | Orto Botanico Forestale dell'Abetone, Italy |
| ***C. tophaceus*** | IBF20190145 | 22.10.2019 | Stabielle, Bedonia, Italy |
| ***C. xanthophyllus*** | IBF20190109 | 20.10.2019 | Oasi Ghirardi, Bedonia, Italy |
| ***C. trivialis*** | IBF20170555 | 14.10.2017 | Pian di Carniglia, Bedonia |

#### 2.2 Mycochemical Fingerprint of Extracts

##### 2.2.1 UV-Vis Spectroscopic Investigations

Each fungal extract (m = 0.5 mg) was solved in methanol (V = 1 mL). After vortexing, the UV-Vis absorption spectra of the solutions were measured ($\lambda$ = 200 – 800 nm) in a glass cuvette (Hellma type 100 – Macro cells, quartz glass, light pass l = 10 mm).

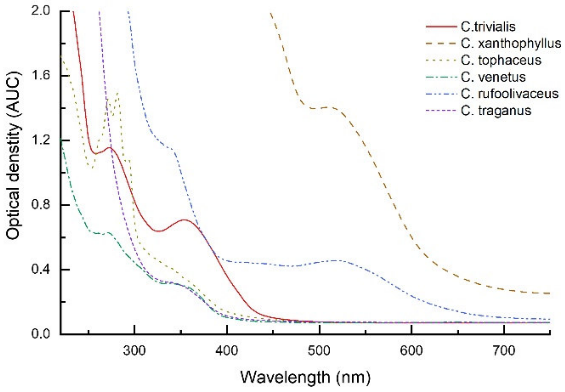


**Figure S 7.** UV-VIS absorption spectra of the investigated extracts (0.5 mg/ml, Methanol).

##### 2.2.2 HPLC-DAD-ELSD/FLD/MS Analysis

All prepared extracts were solved in DMSO (c = 1 mg/ml) and filtered through cotton wool prior to analysis. The setting of the HPLC-DAD was kept constant (Table S1), while the parameters for the other hyphenated detectors (i.e., ELSD, FLD, MS) were changed individually. The mobile phases equaled water (A) and acidified acetonitrile (ACN + 0.1% formic acid (FA), B). As stationary phase, a Synergi MAX-RP (80Å, 4 µm, 150 x 4.60 mm) column was used. The injection volume was V = 5 µl. The flow rate was set to Q = 0.5 mL/min. Column temperature was set to T = 35°C.

**Table S2.** Solvent gradient used for the HPLC-DAD-X analysis

| Time [min] | Solvent A [%] $H_2O$ | Solvent B [%] ACN + 0.1% FA |
|---|---|---|
| 0.00 | 90 | 10.0 |
| 40.0 | 10 | 90.0 |
| 50.0 | 2.0 | 98.0 |
| 51.0 | 90.0 | 10.0 |
| 55.0 | 90.0 | 10.0 |

As second detector, a fluorescence detector (FLD, Figure S8A, $\lambda_{exc}$ = 455 nm, $\lambda_{em}$ = 530 nm), an evaporative light scattering detector (ELSD, Figure S8B), or a mass spectrometer (MS, data not shown) were employed. In Figure S9 the results of the DAD measurement at $\lambda$ = 254 nm are depicted. Figure S10 displays the results of the DAD measurement at $\lambda$ = 428, 478, 528, and 598 nm.

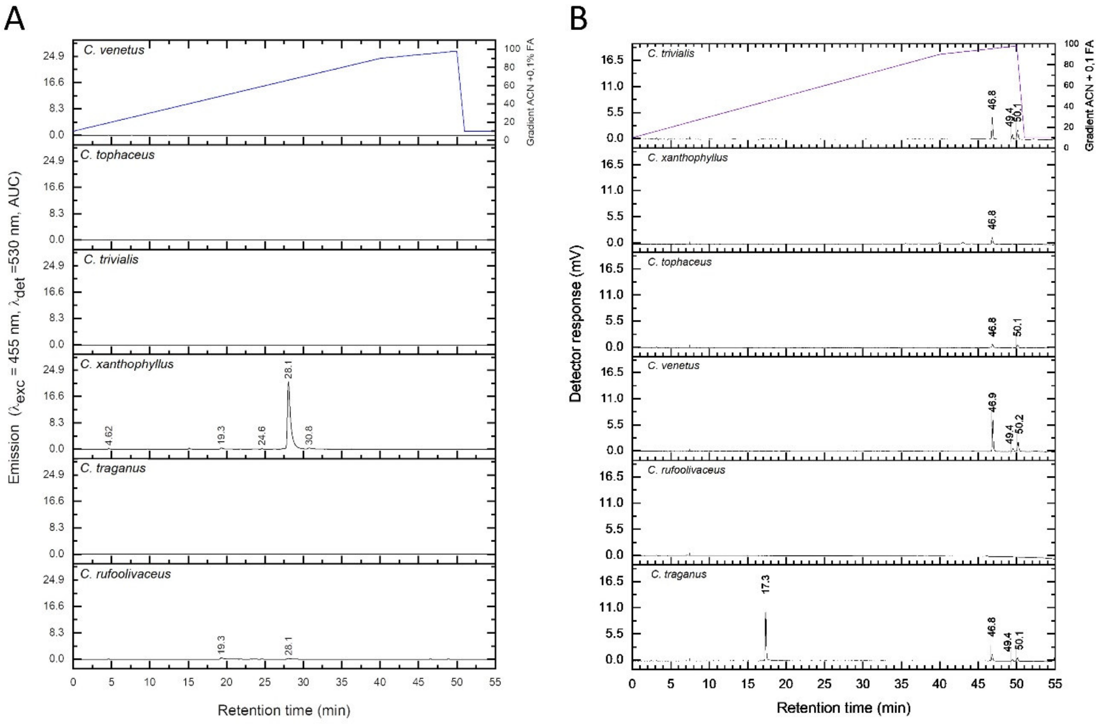


**Figure S8** HPLC-FLD (A) and HPLC-ELSD (B) analysis of the six different extracts. The blue graph represent percent of ACN (+0.1% FA) in the mobile phase (H2O/(ACN+0.1%FA)).

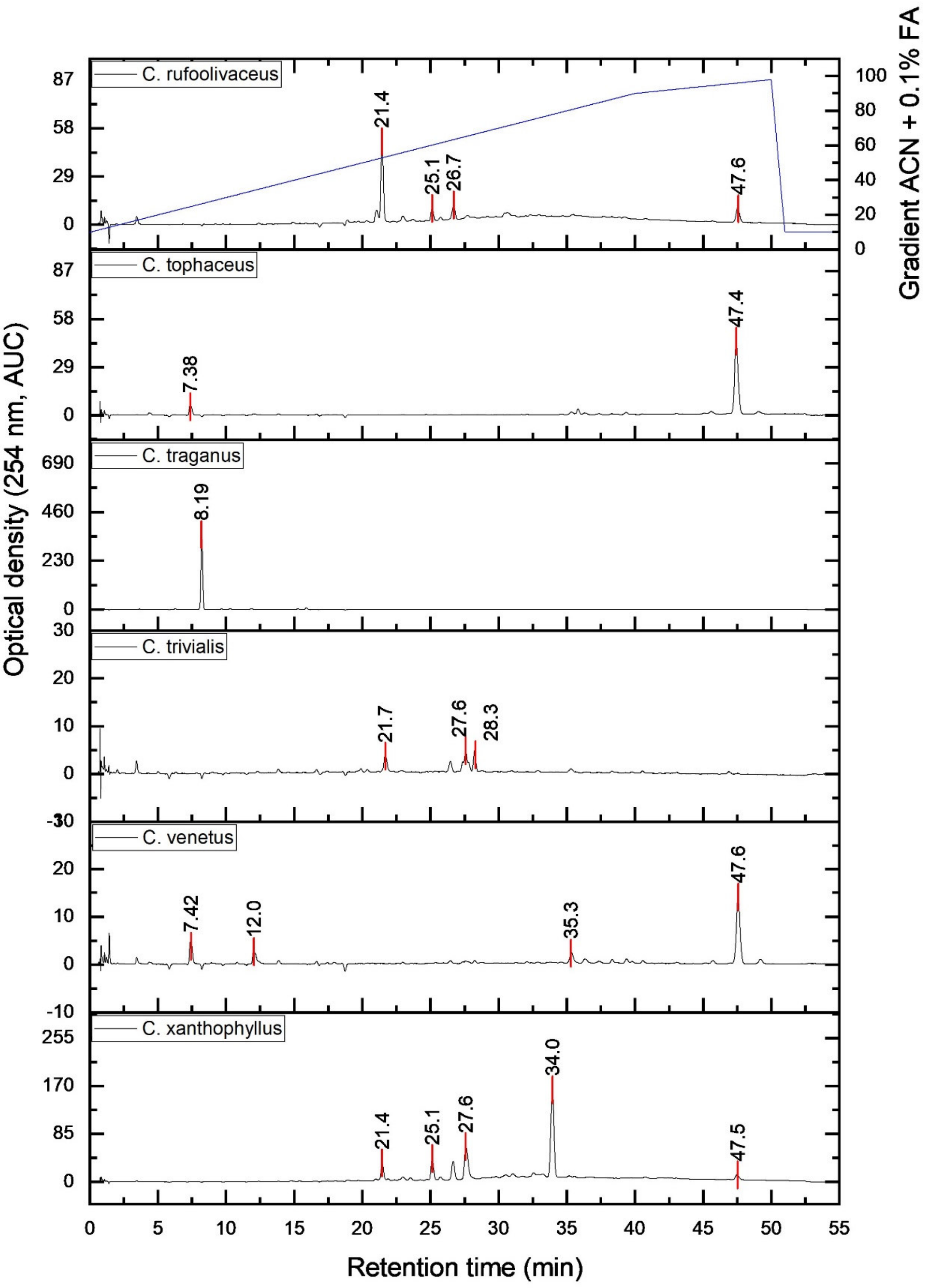


**Figure S9**. HPLC-DAD analysis of the six different extracts (c = 10 mg/ml) detected at λ = 254 nm. The blue graph represent percent of ACN (+0.1% FA) in the mobile phase ($H_2O$/(ACN+0.1%FA)).

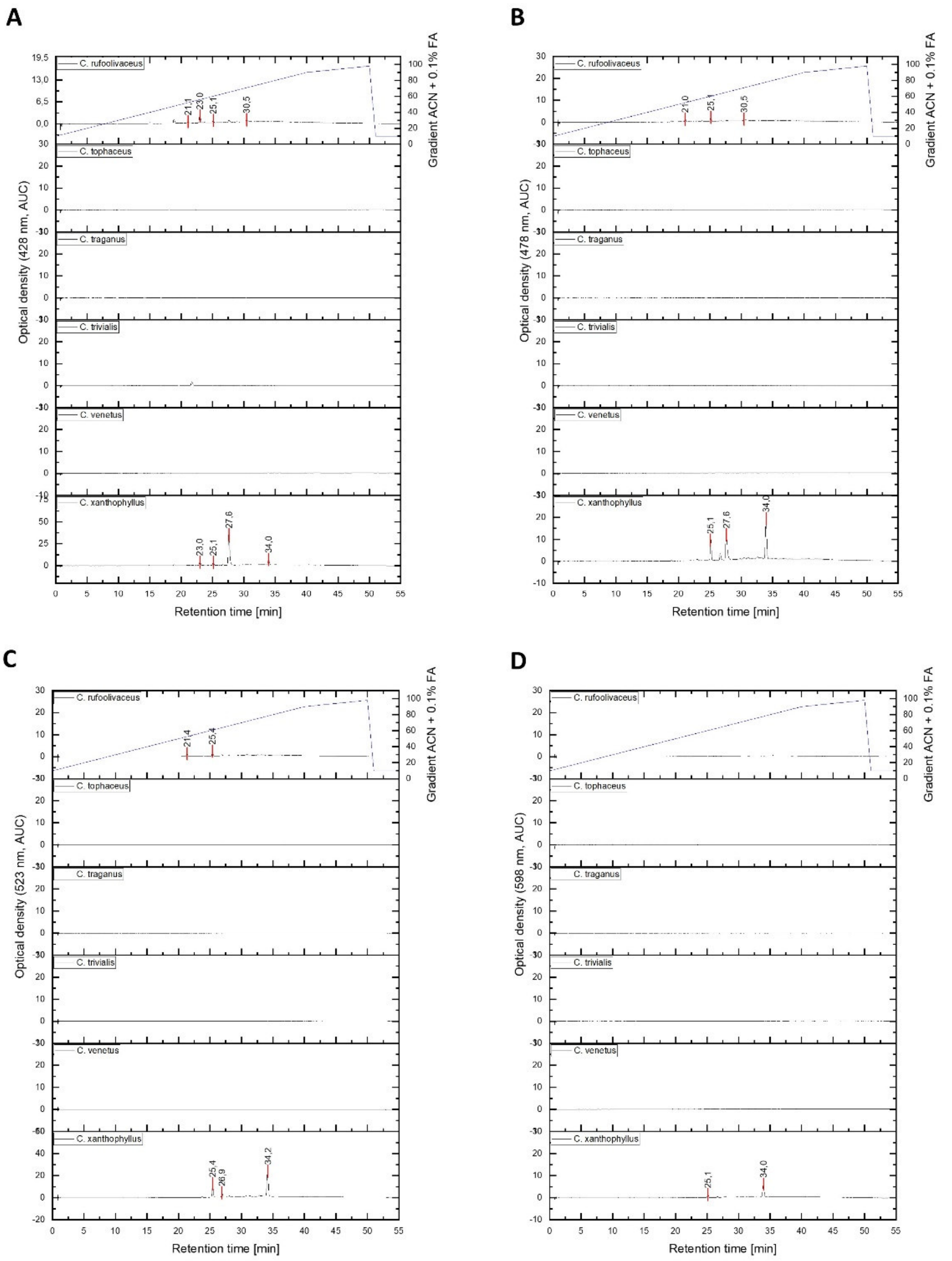


**Figure S10**. HPLC-DAD analysis of the six different extracts (c = 10 mg/ml). Detection wavelength A) 428 nm, B) λ = 478 nm, C) λ = 523 nm, D) λ = 598 nm. The blue graph represent percent of ACN (+0.1% FA) in the mobile phase ($H_2O$/(ACN+0.1%FA)).

#### 2.2.3 General mycochemical analysis

The recorded FLD chromatograms (excitation wavelength of λ = 455 nm and emission wavelength of λ = 530 nm) of most species showed no significant peak. For *C. xanthophyllus*, however, one major peak ($t_r$ = 28.1 min) was detected and correlated to Peak 4 (Figure S10A, $t_{ret}$ = 27.6 min, Table S3). In traces this peak was also detected in *C. rufoolivaceus*. Furthermore, minor peaks were detected at $t_r$ = 19.3 min for both extracts. In addition, for *C. xanthophyllus* two additional peaks at $t_r$ = 24.6 and 30.8 min were observed.

The peak in the ELSD chromatogram (Fig S2B) at $t_r$ = 46.8 min could be detected in all six extracts. It corresponded to a m/z of 279 $[M-H]^-$ in the HPLC-MS experiment and was putatively assigned as linoleic acid. The latter was found as major fatty acid in extracts of the related *Cortinarius magellanicus* (Toledo et al., 2016). Next to this peak, some minor peaks were detected with a similar retention time ($t_r$ = 49.2 and 50.1 min) in several extracts. Furthermore, only in *C. traganus* a peak was detected by the ELSD detector with a retention time of $t_r$ = 17.3 min, indicating a more polar compound.

##### 2.2.4 Mycochemical analysis of *C. xanthophyllus*

The extract of *C. xanthophyllus* was characterized in the HPLC-DAD chromatogram by one major and four minor peaks which absorb at 254 nm. In detail, the major peak is peak **5** with a retention time of $t_r$ = 34.2 min. The minors are peak **1-4** with a retention times of $t_r$ = 21.7, 25.4, 26.9, and 28.0 minutes, respectively. The UV-Vis spectrum of each peak is depicted in Figure S12 and revealed that two anthraquinones with a $\lambda_{max}$ < 500 nm and three anthraquinones with a $\lambda_{max}$ > 500 nm contribute to the overall pigmentation of *C. xanthophyllus*. By the means of a HPLC-DAD-MS experiment the molecular weight of each peak was determined (Table S3).

**Table S3.** Tentative annotation of the pigments from *C. xanthophyllus*

| | $t_r$ [min] | Intensity ($\lambda$=254nm) | $[M-H]^-$ | $[M+H]^+$ | $\lambda_{max}$ [nm] | Proposed Chemical Formula | Suggested Compound |
|---|---|---|---|---|---|---|---|
| 1 | 21.7 | Minor | 645.6 (100%) | 243.1 (26%), 261.1 (100%), 543.2 (15%) | 210 | - | n.d. |
| 2 | 25.4 | Minor | 555.3 (100%) | 557.2 (100%) | 216, 300, 514 | $C_{32}H_{28}O_9$ | Rufoolivacin A |
| 3 | 26.9 | Minor | - | 498.4 (23%) 519.2 (16%) 557.2 (100%) | 298, 335, 493 | $C_{32}H_{28}O_9$ | Rufoolivacin C |
| 4 | 28.0 | Minor | 608.8 (25%) 624.9 (100%) | 285.1 (7%) 377.4 (6%) 474.5 (10%) 543.2 (100%) 585.2 (23%) | 222, 270, 288, 436 | $C_{16}H_{12}O_5$ | Physcion/Parietin |
| 5 | 34.2 | Major | - | 256.3 (9%) 409.4 (10%) 459.4 (10%) 501.4 (10%) 556.2 (100%) | 228, 258, 335, 525 | - | n.d. |

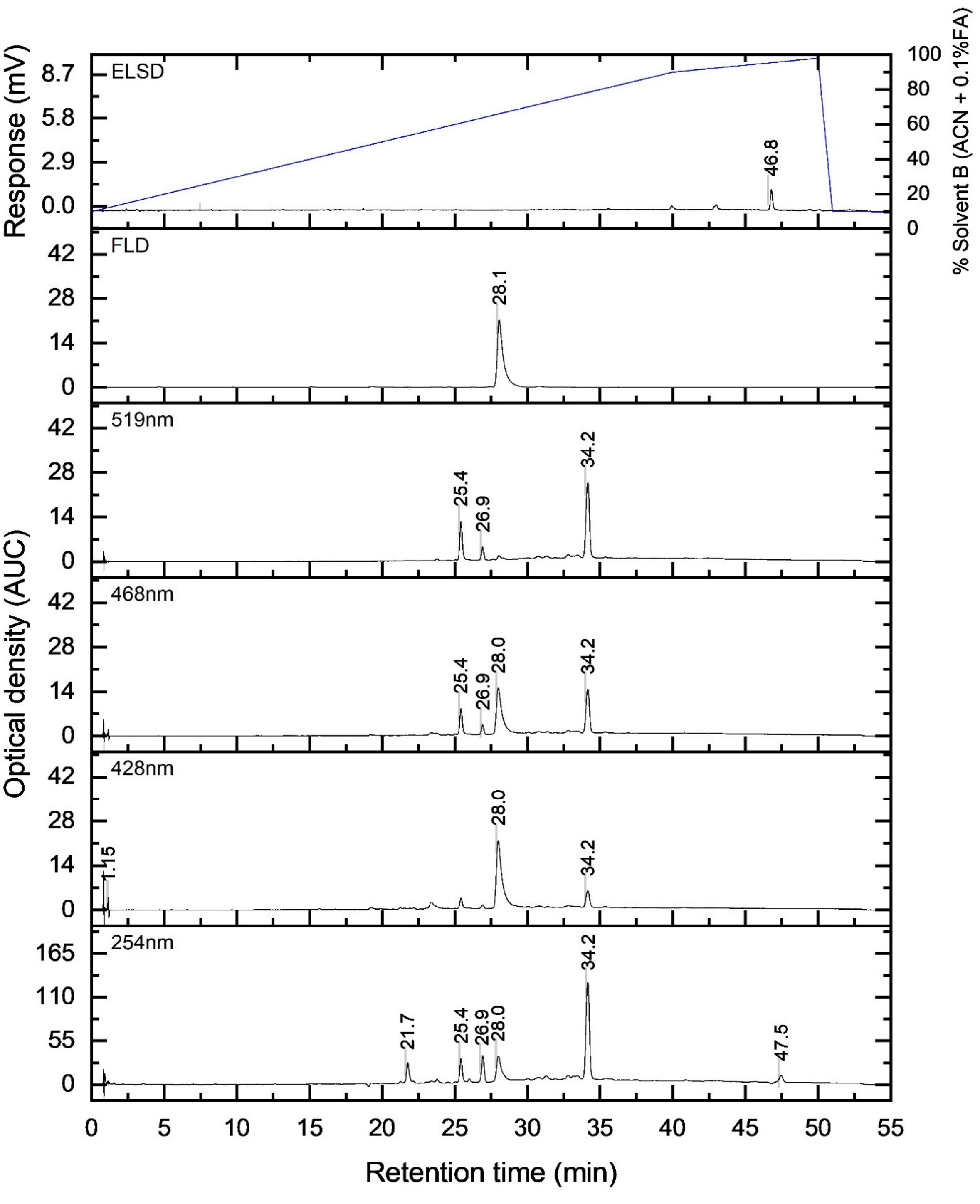


**Figure S11.** HPLC-DAD-X Chromatograms of the *C. xanthophyllus* extract (1 mg/ml, DMSO): From the bottom to the top the chromatograms detected at $\lambda = 254$ nm, 428 nm, 468 nm, 519 nm are depicted as well as the FLD ($\lambda_{exc} = 455$ nm, $\lambda_{det} = 530$ nm) and ELSD chromatogram.

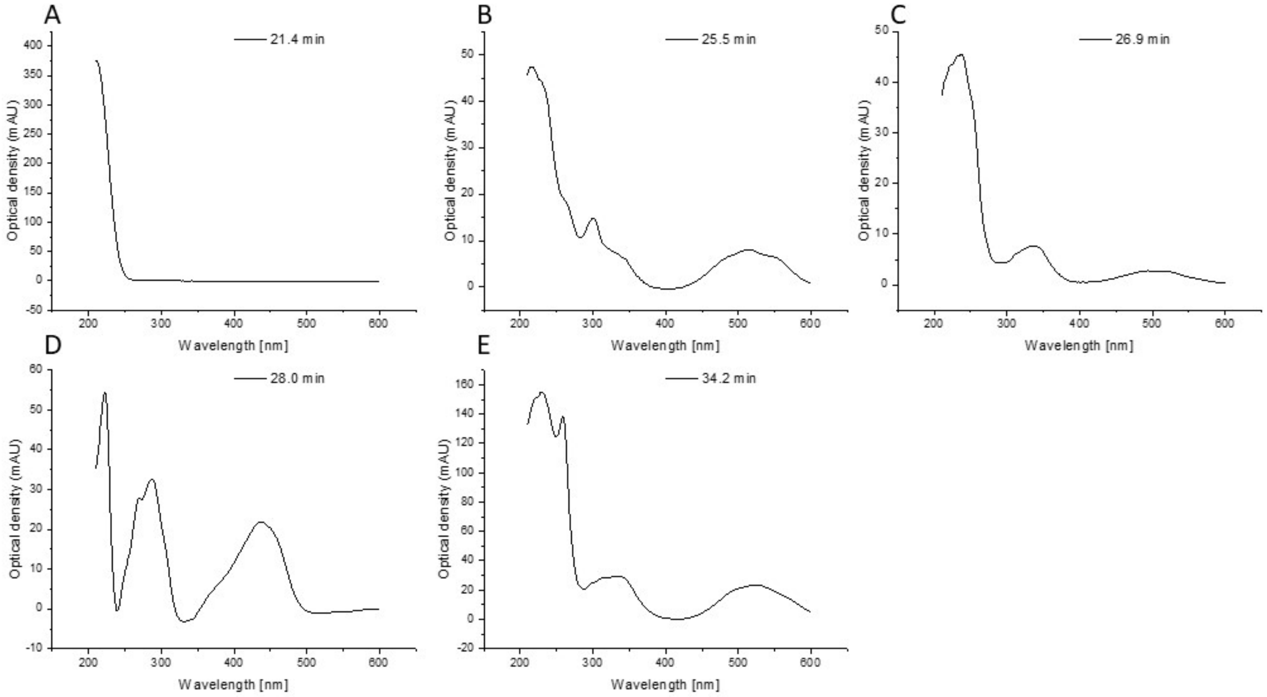


**Figure S12:** UV-Vis spectra of the major and minor peaks of *C. xanthophyllus* extracted from the HPLC-DAD experiment. Peaks are sorted according to their retention time, A) $t_r$ = 21.4 min, B) $t_r$ = 25.5 min, C) $t_r$ = 29.9 min, D) $t_r$ = 28.0 min, E) $t_r$ = 34.2 min.

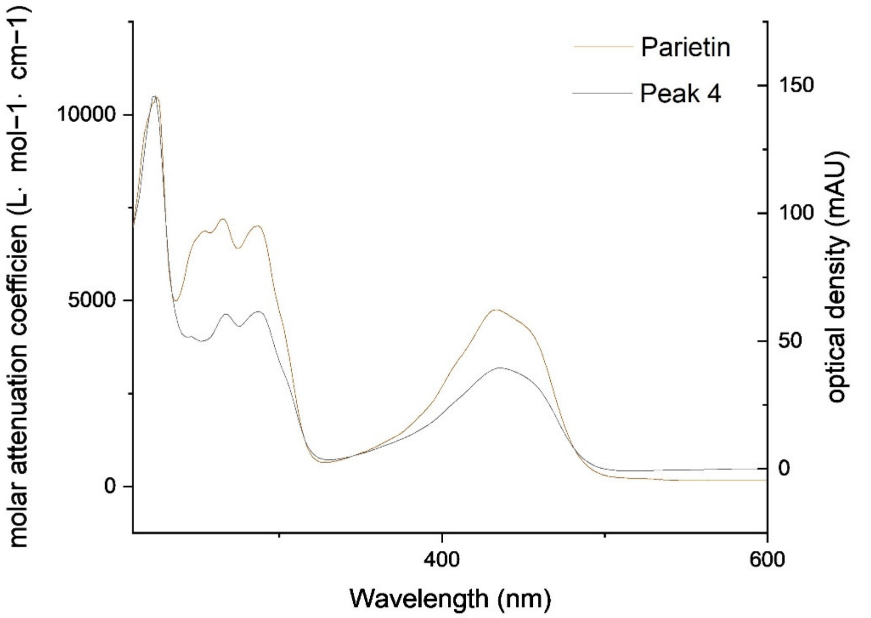


**Figure S13:** UV-VIS overlay of parietin (reference compound) and peak **4** of the *C. xanthophyllus* extract. The minor differences can be attributed to the different solvents used (i.e., MeOH and ACN+0.1%FA, respectively).

### 3 (Photo)antimicrobial evaluation of the *C. xanthophyllus* extract

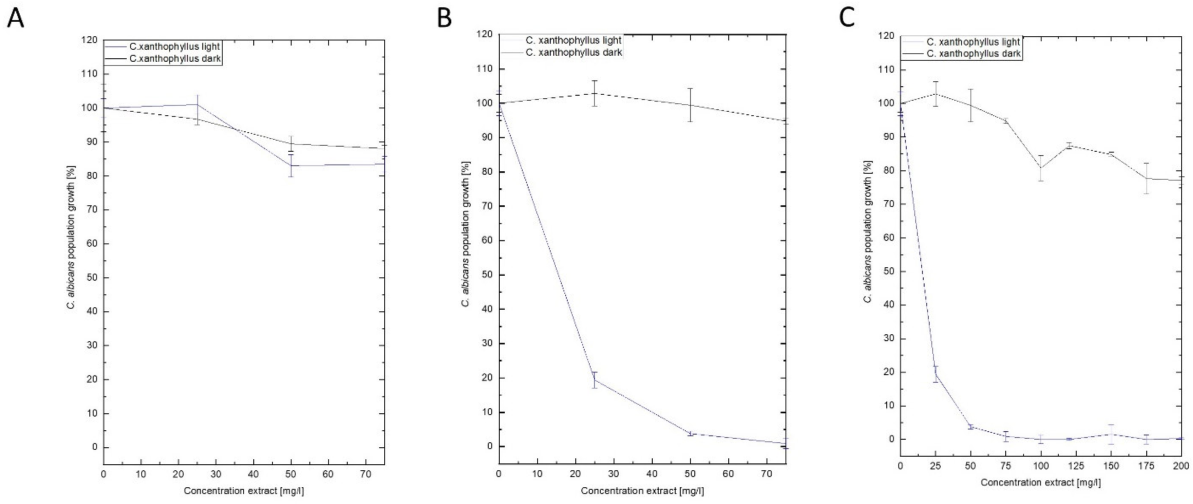


**Figure S14.** (Photo)antimicrobial action of *C. xanthophyllus* extract against *C. albicans* under blue light irradiation ($\lambda$ = 468 nm, $H$ = 30 J/cm$^2$) in dependency of the pre-incubation time. A) PI = 10 min, B) PI = 60 min, C) PI = 60 min and an extended concentration range showing a large therapeutic window.
